# A conserved ubiquitin- and ESCRT-dependent pathway to regulate human lysosomal membrane proteins

**DOI:** 10.1101/2020.11.18.389296

**Authors:** Weichao Zhang, Xi Yang, Liang Chen, Yun-Yu Liu, Varsha Venkatarangan, Lucas Reist, Phyllis Hanson, Haoxing Xu, Yanzhuang Wang, Ming Li

## Abstract

The lysosome is an essential organelle to recycle cellular materials and maintain nutrient homeostasis, but the mechanism to down-regulate lysosomal membrane proteins is poorly understood. In this study, we developed a cycloheximide chase assay to measure the half-lives of ~30 human lysosomal membrane proteins, and identified RNF152 as a short-lived protein. The degradation of RNF152 depends on ubiquitin and the endosomal sorting complexes required for transport (ESCRT) machinery. Ubiquitinated RNF152 is sorted and internalized by the ESCRT machinery into the lysosomal lumen for degradation. Strikingly, when expressed in budding yeast, human RNF152 is also degraded by the vacuole (yeast lysosome) in an ESCRT-dependent manner. Thus, our study uncovered a conserved mechanism to down-regulate lysosome membrane proteins.

## Introduction

As an essential organelle, the lysosome is responsible for various cellular processes, including protein turnover and recycling, energy metabolism, intracellular signaling, and nutrient storage (Lawrence and Zoncu, 2019). The lysosome membrane contains hundreds of transmembrane proteins, many of which are transporters and channels that shuttle cargoes (ions, amino acids, cholesterol, etc.) across the lysosomal membrane (Schröder et al., 2007; Chapel et al., 2013; Bissa et al., 2016; Wyant et al., 2018). Malfunction of these lysosomal membrane proteins (LMPs) can give rise to inherited genetic disorders called lysosomal storage diseases (LSDs). Without treatment, almost all types of LSD patients will develop severe neurodegeneration in the central nervous system. Furthermore, growing evidence suggests that mutations in LMPs and lysosome dysfunction are associated with age-related neurodegeneration diseases such as Alzheimer’s disease, frontotemporal dementia, and Parkinson’s disease (Nixon et al., 2006; Amick and Ferguson et al., 2017; Cook et al., 2012). As we age, the lysosome membrane gradually accumulates damaged proteins and loses its activity, which dampens the cell’s ability to remove pathogenic protein aggregates and damaged organelles, eventually leading to cell death and inflammation (Carmona-Gutierrez et al., 2016; Cheon et al., 2019; Yambire et al., 2019). Strategies to maintain the lysosome membrane integrity during aging will likely delay the onset of neurodegenerative symptoms.

Given the physiological importance and clinical implications of LMPs, we wonder how LMPs are regulated and quality controlled. At the organelle level, if the lysosomal membrane is mildly damaged by insults like lysosomotropic compounds or iron-dependent oxidative stress, the endosomal sorting complexes required for transport (ESCRT) machinery can be recruited to the lysosome surface to repair the membrane (Mittal et al., 2018; Radulovic et al., 2018; Skowyra et al., 2018). If the lysosome damage is too severe to be repaired, ruptured lysosomes will be sequestered and degraded by selective autophagy, a process termed as lysophagy (Hung et al., 2013; Maejima et al., 2013).

At the protein level, removal of specific membrane proteins from the lysosome surface is essential for adjusting its membrane protein composition in response to environmental cues. However, very little is known about the mechanism underlying this process. A process like lysophagy, which engulfs whole lysosomes, could not possibly turn over a few proteins selectively. This leads us to ask important questions as to how the human lysosome selectively downregulates its membrane proteins and what machinery might be involved in the process.

In this study, we screened ~30 human LMPs using a cycloheximide chase assay and identified a few candidates with short half-lives. Among those candidates, we focused on RNF152 (a lysosome membrane-anchored E3 ligase) as cargo to examine the possible mechanisms of LMP turnover in human cells. We uncovered that the degradation of RNF152 is both ubiquitination- and lysosome-dependent. Further, we provided evidence that the conserved ESCRT machinery plays an important role in the internalization of RNF152. Collectively, our work suggests that the ubiquitin- and ESCRT-dependent degradation pathway is a conserved mechanism to downregulate LMPs in both yeast and human cells.

## RESULTS

### Cycloheximide (CHX) Chase Screen to Determine the Half-lives of LMPs

To study the regulation and quality control of lysosomal membrane proteins (LMPs) in human cells, we urgently needed a substrate with fast turnover. To this end, we collected more than 30 GFP- or mCherry-tagged human lysosome membrane proteins. These proteins were identified as lysosomal by either microscopy or Mass Spectrometry studies (Schröder et al., 2007; Chapel et al., 2013; Schwake et al., 2013; Bissa et al., 2016; Wyant et al., 2018). We transiently transfected HEK293 cells with plasmids encoding these membrane proteins and determined their half-lives using the cycloheximide chase assay. Cycloheximide is a protein synthesis inhibitor that allows us to measure the half-lives of the pre-existing protein population (Kao et al., 2015). Here, we present some examples of the cycloheximide chase screen (Figure 1A-B, Figure S1, and Table 1). While most LMPs are relatively stable (examples: TMEM175, PQLC2, CTNS, CLCN7, LAMP2, and TMEM192), a few LMPs have half-lives less than 9 hours (e.g., LAPTM4A, RNF152, and OCA2) (Figure 1A-B). For LAPTM4A, RNF152, and OCA2, we confirmed their lysosome localizations by co-staining with the lysosome-associated membrane protein 2 (LAMP2) (Figure 1C). Of note, a small fraction of GFP-RNF152 puncta do not colocalize with LAMP2 (white arrows in Figure 1C). This observation is consistent with a recent study indicating that RNF152 could also localize to endosomes (Xiong et al., 2020).

**Figure 1:**
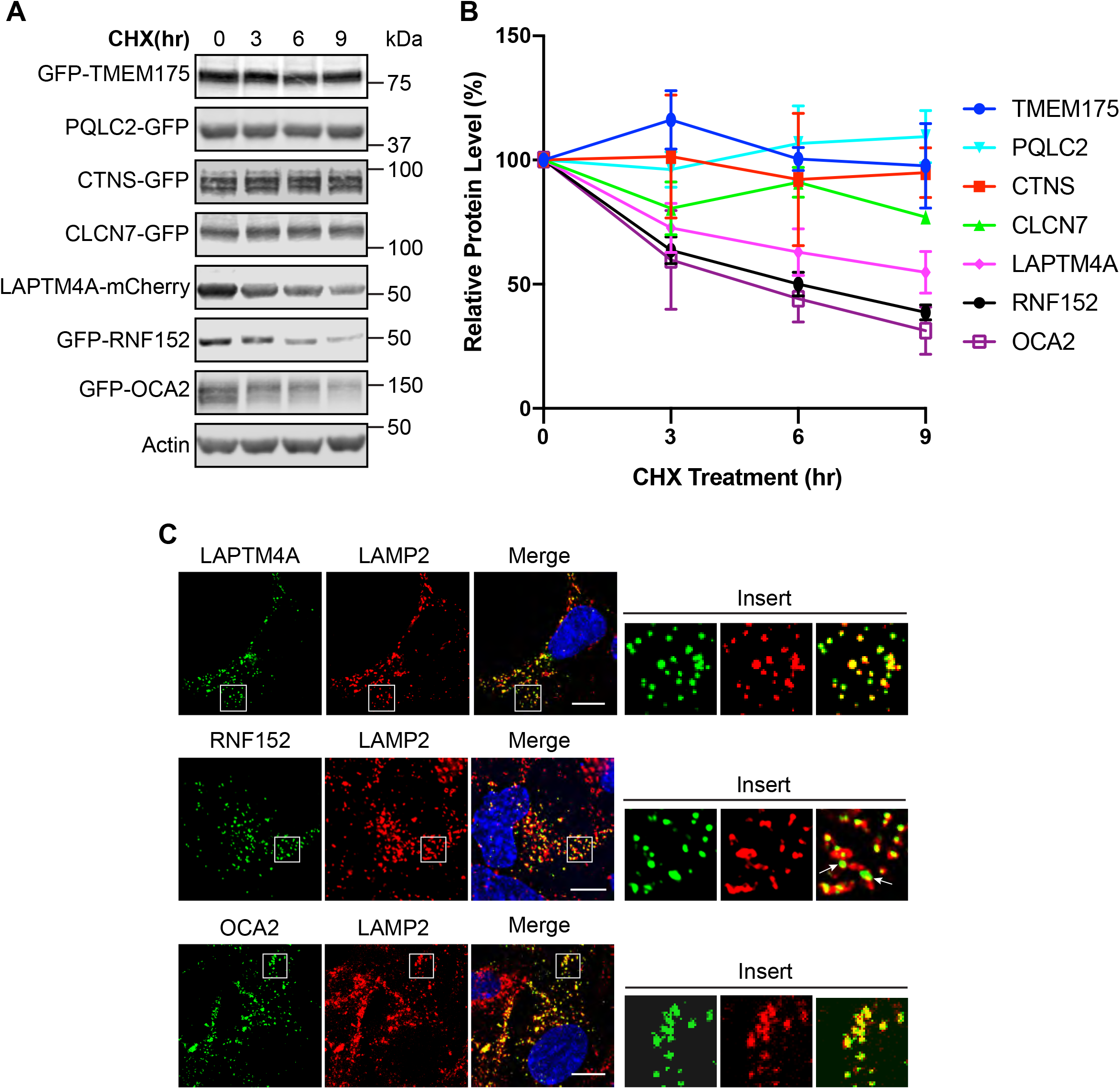
Cycloheximide chase assay to screen for LMPs with short half-lives. (A) Western blots showing the level of indicated LMPs in cells treated with cycloheximide (CHX) for the indicated time periods. HEK293 cells were transfected with indicated plasmids. CHX chase assay was performed 20-24h post-transfection and detected with either a GFP or mCherry antibody. (B) Quantification of the protein level in A, n=3. (C) Immunofluorescence showing the localization of LAPTM4A-mCherry, GFP-RNF152, and GFP-OCA2 in HEK293 cells costained with a LAMP2 antibody. LAPTM4A, RNF152, and OCA2 are pseudo-colored in green. LAMP2 is pseudo-colored in red. Scale bar=10μm. White arrows indicate GFP-RNF152 puncta that are not colocalized with LAMP2 signals.

Although OCA2 shows the fastest degradation kinetics, it is a melanosome protein that does not exist in normal lysosomes (Sitaram et al., 2009). Studying the degradation of OCA2 in cells other than melanocytes may be physiologically irrelevant. Therefore, we focused on the other fast turnover protein RNF152 to investigate the degradation pathway for lysosome membrane proteins.

### RNF152 is a protein of short half-life

RNF152 is a single pass, Type-II transmembrane protein with 203 amino acids (Figure 2A). It was recently characterized as a lysosome E3 ligase that negatively regulates mTORC1 activity by ubiquitinating RagA (Deng et al., 2015). We tagged RNF152 with GFP at its N-terminus so that GFP is facing the cytosol. The molecular weight of RNF152 is ~23 kDa. After fusing with GFP, the calculated size is around 50kDa.

**Figure 2:**
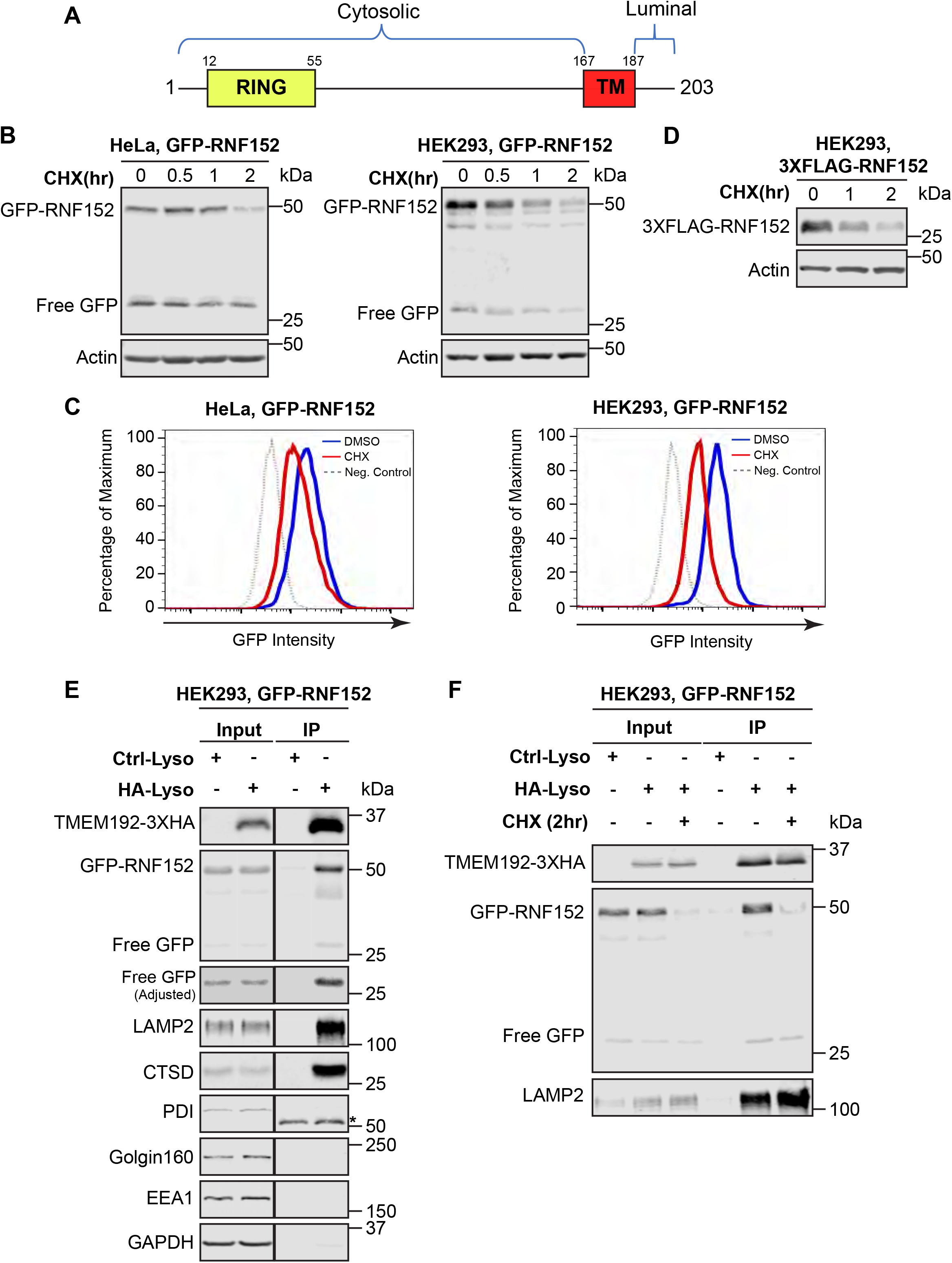
RNF152 has a short half-life. (A) Domain organization of RNF152. (B) Western blots showing the degradation of GFP-RNF152 in HeLa and HEK293 cells. GFP-RNF152 is stably expressed in these cell lines. (C) Flow Cytometry analysis showing GFP intensity in HeLa and HEK293 cells that stably express GFP-RNF152. Cells were incubated with indicated treatments for 2 hours before the Flow Cytometry analysis. Non-infected HeLa and HEK293 cells served as negative controls. (D) Western blots showing the degradation of 3XFLAG-RNF152 in HEK293 cells. 3XFLAG-RNF152 is stably expressed. (E) Representative Western blots showing both full-length GFP-RNF152 (50kDa) and free GFP (27kDa) are enriched after Lyso-IP. LAMP2 (lysosomal membrane), CTSD (lysosomal lumen), PDI (ER), Golgi160 (Golgi), EEA1 (endosome), and GAPDH (cytosol) were also immunoblotted. Ctrl-Lyso: cells stably expressing TMEM192-2XFLAG; HA-Lyso: cells stable expressing TMEM192-3HA. Both cell lines also stably express GFP-RNF152. Asterisk: mouse heavy chain. (F) Representative Western blots showing the degradation of lysosome localized GFP-RNF152 (50kDa) after two-hour cycloheximide treatment. LAMP2 serves as an internal control.

Our original screen was conducted with transient transfection. To avoid variations of GFP-RNF152 expression in cells, we generated cell lines (HEK293 and HeLa) that stably express GFP-RNF152 by lentivirus infection. We treated these stable cell lines with cycloheximide and confirmed that the half-life of GFP-RNF152 is about 1 hour in HEK293 cells and 2 hours in HeLa cells, respectively (Figure 2B). Besides Western blots, we also used Flow Cytometry analysis to confirm the degradation of GFP-RNF152 in these cell lines. As shown in Figure 2C, the GFP intensity in both HEK293 and Hela cell lines was significantly reduced after 2 hours of cycloheximide treatment. Lastly, to rule out the possibility that the degradation of GFP-RNF152 is due to the GFP fusion, which is larger than RNF152, we replaced GFP with a much smaller 3xFlag tag (23 amino acids). As shown in Figure 2D, 3xFlag-RNF152 was also quickly degraded in HEK293 cells.

Interestingly, we observed a 27kDa band on the GFP blot that is about the size of free GFP in the cycloheximide chase assay (Figure 2B). Because GFP is fused to the cytosolic side of RNF152 (Figure 2A), there are two possibilities to generate the 27 kDa band. First, GFP-RNF152 may be internalized into the lysosome lumen for degradation, and the free GFP is the by-product in the lysosome lumen. Second, free GFP may be cleaved off RNF152 due to the instability of the fusion protein, in this case free GFP would be localized in the cytosol. To differentiate between these two possibilities, we adopted the Lyso-IP experiment developed by the Sabatini group (Abu-Remaileh et al., 2017). The Lyso-IP assay uses a 3xHA-tagged lysosome-specific membrane protein TMEM192 as the bait to pull down intact lysosomes. As expected, the full-length GFP-RNF152 and other lysosomal proteins such as LAMP2 and cathepsin D (CTSD) were enriched by Lyso-IP. In contrast, PDI (ER), Golgin160 (Golgi), EEA1 (endosomes), and GAPDH (cytosol) were not enriched (Figure 2D). Importantly, we found that the 27 kDa GFP band was also enriched by Lyso-IP, which indicates that the free GFP is inside the lysosome lumen. These data suggest that RNF152 may be internalized and degraded in the lysosome.

Because a fraction of RNF152 localizes to the endosome (white arrows in Figure 1C, Xiong et al., 2020), it is possible that the observed degradation is due to the endosomal degradation of RNF152. To ensure that the lysosome population of RNF152 is fast-degrading, we performed the Lyso-IP experiment using cells collected before and after 2 hours of CHX treatment. Our result confirmed that the lysosome population of GFP-RNF152 is quickly turned over, while LAMP2 is very stable on the lysosome (Figure 2E).

### RNF152 degradation is ubiquitination-dependent

Protein ubiquitination is the prerequisite for degradation in eukaryotic cells. Given that RNF152 has a short half-life, we hypothesized that RNF152 might be ubiquitinated continuously in the cell. The full-length GFP-RNF152 appears as a 50 kDa band on the Western blots. Interestingly, we also observed a faint 60 kDa band that might be the mono-ubiquitinated form of GFP-RNF152 (Figure 3A, left panel). After longer exposure, a high-molecular-weight smear appeared on top of the full-length GFP-RNF152 (Figure 3A, right panel). Could this smear be the poly-ubiquitinated GFP-RNF152? To answer this, we transfected HA-tagged ubiquitin into cells that stably express either GFP-RNF152 or free GFP control, followed by immunoprecipitation using the GFP-TRAP nanobody. Indeed, HA-ubiquitin was incorporated into the high-molecular-weight smear of GFP-RNF152, but not the GFP control (Figure 3B). This result confirmed that a significant portion of GFP-RNF152 is constantly poly-ubiquitinated.

**Figure 3:**
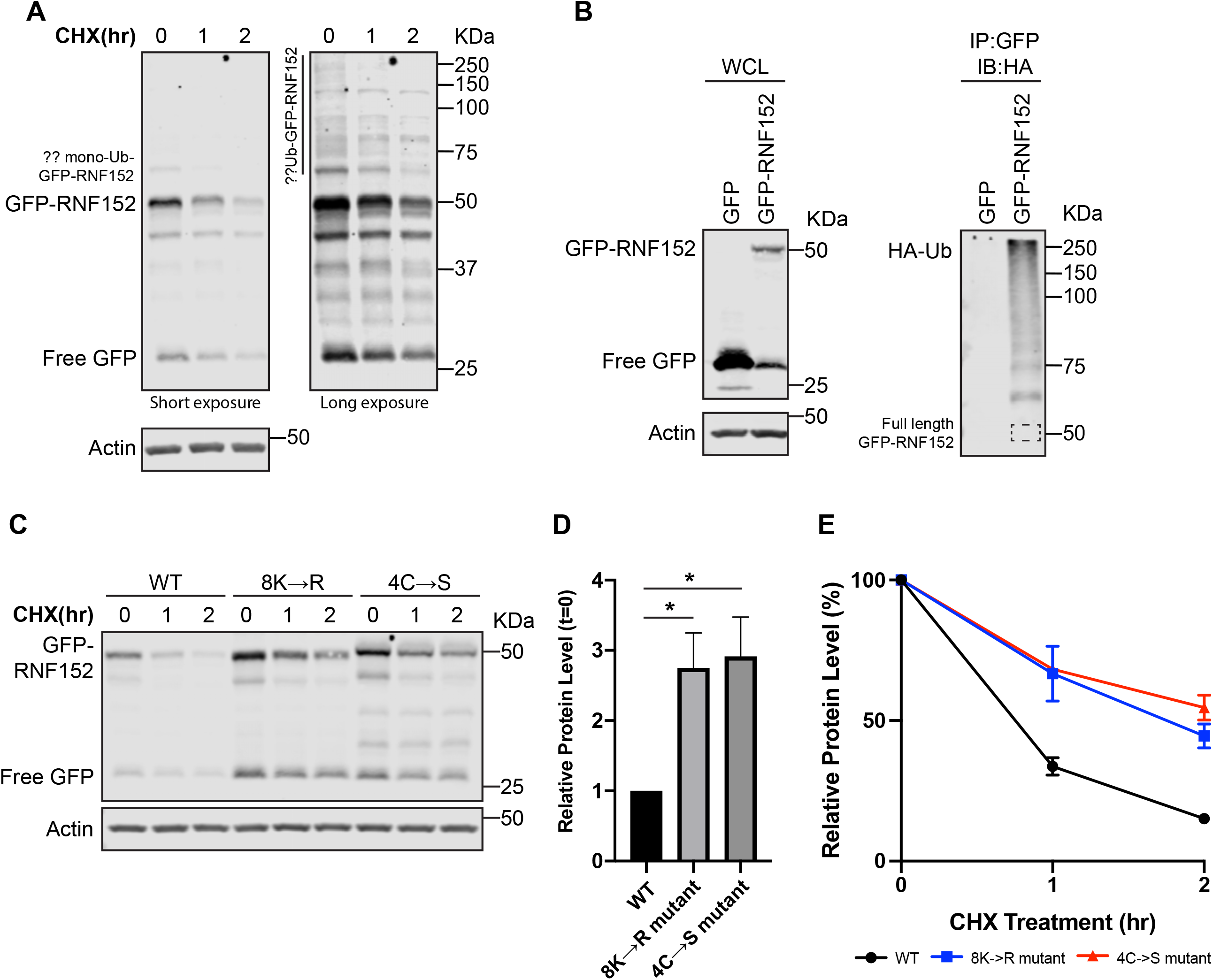
RNF152 degradation is ubiquitin-dependent. (A) Left panel: representative Western blots showing GFP-RNF152 degradation in HEK293 stable cell line. A faint band at 60kDa might be the mono-ubiquitinated form of GFP-RNF152. Right panel: long exposure of the left panel. A high-molecular-weight smear appears on top of the GFP-RNF152, which might be poly-ubiquitinated GFP-RNF152. (B) Left panel: whole cell lysate (WCL) of HEK293 cells that stably express either GFP or GFP-RNF152 and transfected with HA-Ub. Right panel: GFP or GFP-RNF152 were immunoprecipitated with GFP-TRAP nanobody and immunoblotted with an HA antibody. A representative Western blot shows that GFP-RNF152 but not free GFP is poly-ubiquitinated. The dotted box represents the size of full-length GFP-RNF152 at 50kDa. (C) Representative Western blots showing degradation kinetics of wildtype (WT) GFP-RNF152, 8K→R and 4C→S mutants in HEK293 cells that stably express each construct. (D) Quantification of the protein level of WT GFP-RNF152 and its mutants at the steady-state (t=0h in C), n=3. * p<0.05. (E) Quantification of the protein level of WT GFP-RNF152 and its mutants in cycloheximide chase assay (all time points in C), n=3.

To test if poly-ubiquitination of GFP-RNF152 is important for its degradation, we mutated all eight lysines in the cytosolic domain of RNF152 to arginine (8K→R). As shown in Figure 3C-3D, the 8K→R mutation increased the steady-state level of GFP-RNF152_8K→R_ (2.7 fold at time 0). Additionally, the degradation kinetics of the mutant was also significantly slowed down (Figure 3E).

Because RNF152 is a RING domain-containing E3 ligase, we hypothesized that the degradation of GFP-RNF152 might be at least partially due to its auto-ubiquitination. To test this hypothesis, we mutated four cysteines in the RING finger motif to serines (4C→S mutant) to disrupt its E3 ligase activity (Deng et al., 2015). Similar to the 8K→R mutant, the protein level of GFP-RNF152_4C→S_ at the steady-state was increased by 2.8-fold (t=0h, Figure 3C-3D), and the degradation kinetics was significantly slower (Figure 3E).

Taken all together, we concluded that the degradation of GFP-RNF152 is a ubiquitin-dependent process. The short half-life of GFP-RNF152 is likely due to its auto-ubiquitination.

### RNF152 is mainly degraded by the lysosome

In eukaryotic cells, there are two primary mechanisms responsible for breaking down proteins: proteasome-dependent and lysosome-dependent proteolysis. The accumulation of free GFP inside the lysosome suggests that RNF152 is degraded by the lysosome. To test this hypothesis further, we treated the cells that stably express GFP-RNF152 with either lysosome V-ATPase inhibitor Bafilomycin A1 (BafA1) or proteasome inhibitor MG132. Inhibition of V-ATPase results in a rapid neutralization of lysosome pH, leading to the Inhibition of luminal protease activities. After 6 hours of BafA1 treatment, the steady-state protein level of full-length GFP-RNF152 increased by two-fold (Figure 4A-B). In contrast, MG132 treatment did not lead to a significant change in the GFP-RNF152 protein level, while the double treatment did not further increase the steady-state RNF152 level than BarfA1 alone.

**Figure 4:**
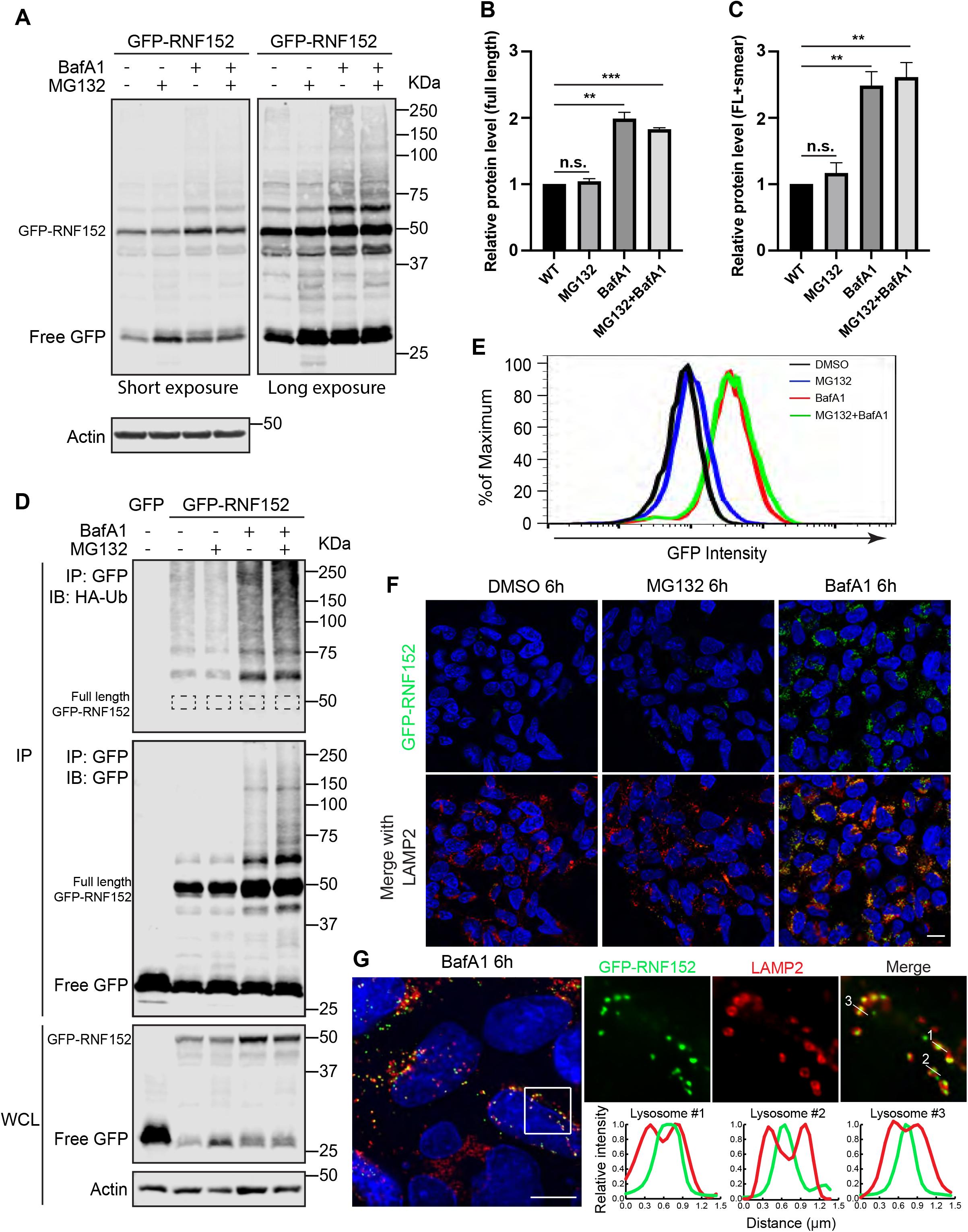
RNF152 is mainly degraded by the lysosome. (A) Left panel: representative Western blots showing GFP-RNF152 protein level after MG132 (50 μM), BafA1 (400 nM), and double treatment for 6 hours. Right panel: long exposure of the left panel. (B) Quantification of the full-length GFP-RNF152 protein level at 50 kDa in A, n=3. n.s. not significant. ** p <0.01, *** p <0.001. (C) Quantification of the total GFP-RNF152 protein level, including full-length at 50kDa plus all high-molecular-weight smear in A, n=3. n.s. not significant. ** p <0.01. (D) GFP or GFP-RNF152 were immunoprecipitated with GFP-TRAP nanobody and immunoblotted with antibodies against HA tag (Top panel) and GFP (Mid panel). Dotted boxes represent the size of full-length GFP-RNF152. Bottom panel: Western blots showing whole cell lysate (WCL). Cells stably expressing free GFP served as a negative control. All cell lines were transfected with HA-Ub plasmids. Cells were treated with MG132 (50 μM), BafA1 (400 nM), or MG132+BafA1 for 6h. (E) Flow Cytometry analysis showing GFP intensity of HEK293 cells that stably express GFP-RNF152 after 6 hours of indicated treatment. (F) Immunofluorescence of LAMP2 with GFP-RNF152 in HEK293 cells that stably express GFP-RNF152 after indicated treatment. Scale bar=10 μm. (G) Immunofluorescence of LAMP2 with GFP-RNF152 in HEK293 cells that stably express GFP-RNF152 after 6 hours of BafA1 (400 nM) treatment. Line scanning analysis was performed on three lysosomal structures. Scale bar=10 μm.

After extended exposure, we also observed an accumulation of high-molecular-weight smear in the BafA1-treated and the double treatment groups (Figure 4A and 4C). To verify that BafA1 treatment leads to the accumulation of poly-ubiquitinated GFP-RNF152, we co-expressed HA-tagged ubiquitin with GFP-RNF152 and performed immunoprecipitation. We found that the poly-ubiquitinated GFP-RNF152 indeed increased drastically after the BafA1 treatment and BafA1+MG132 double treatment, but not after the MG132 treatment (Figure 4D). These results suggest that the lysosome, but not the proteasome, is the primary organelle to degrade RNF152, which is consistent with the observation that free GFP accumulated inside the lysosome after degradation (Figure 2D).

We further confirmed our findings with Flow Cytometry analysis. As shown in Figure 4E, the GFP intensity only increased slightly after MG132 treatment. In contrast, BafA1 treatment led to a major increase in GFP intensity (7-8-fold increase). BafA1+MG132 double treatment did not lead to a further increase in GFP intensity. Although these Flow Cytometry results consistently suggest that RNF152 degradation is lysosome-dependent, we were surprised by the strong increase (7-8-fold) in GFP intensity after the BafA1 treatment.

To address this concern, we use confocal microscopy to image HEK293 cells that stably express GFP-RNF152 after either DMSO, MG132, or BafA1 treatment. Under the same imaging settings, we found that the GFP intensity after BafA1 treatment was much higher compared with DMSO or MG132 treatment groups. Most of the BafA1-stabilized GFP signals were colocalized with the lysosome marker LAMP2 (Figure 4F). Importantly, when we imaged the BafA1-treated cells with a Delta-Vision fluorescent microscope, which is incorporated with a powerful deconvolution software to enhance the resolution of images, we observed that much of the GFP signal was localized inside the lysosome lumen (line scan in Figure 4G). Similar results were also obtained with a Leica STED super-resolution microscope (data not shown). The luminal GFP signal inside the lysosome strongly supports a model that RNF152 is internalized into the lysosome and degraded by lumenal proteases. Once GFP gets into the lysosome lumen, the fluorescence will be quenched by the low pH (4.5-5). However, BafA1 treatment increases the lumenal pH, and the GFP fluorescence is no longer quenched, which leads to a drastic increase of GFP intensity in the lysosome lumen.

Besides the steady-state protein levels, we also studied the effects of BafA1 and MG132 on the degradation kinetics of GFP-RNF152. As shown in Figure S2A-B, the degradation of GFP-RNF152 was blocked by BafA1 treatment, even when we extended the chase to 6 hours. On the other hand, the proteasome inhibitor MG132 only delays the degradation of GFP-RNF152. Interestingly, we observed an increase of free GFP intensity after both MG132 and BafA1 treatment. MG132 is known to partially inhibit some classes of lysosomal proteases such as Cathepsin A and Cathepsin B (Shirley et al., 2005). The delayed degradation of GFP-RNF152 and increased free GFP signals might be due to inefficient degradation by the lysosomal proteases.

Lastly, we examined the fate of the lysosome-specific population of GFP-RNF152 by lyso-IP. As shown in Figure S2C, the lysosome population of GFP-RNF152 is also stabilized by BafA1 treatment in the CHX chase assay.

Taken all results together, we concluded that RNF152 is mainly degraded through lysosome-dependent proteolysis. For the rest of our study, we will focus on identifying the machinery that can deliver RNF152 into the lysosomal lumen for degradation.

### Macroautophagy machinery and CMA pathways are not involved in the degradation of GFP-RNF152

In mammalian cells, there are four possible mechanisms to deliver intracellular materials into lysosomes for degradation: macroautophagy, microautophagy, ESCRT-dependent formation of intraluminal vesicles at endosomes, and chaperone-mediated autophagy (Schuck, 2020; Schmidt and Teis, 2012; Bejarano and Cuervo, 2010). How is ubiquitinated RNF152 internalized into the lysosome lumen then? Although lysophagy (selective macroautophagy of lysosomes) can deliver an entire damaged lysosome to other healthy lysosomes for degradation (Anding and Baehrecke, 2017), it cannot selectively turnover a particular lysosome membrane protein while leaving others intact. Very recently, Overholzer and colleagues reported that glucose starvation or certain pharmacological drug treatments could trigger a microautophagy process to selectively turnover lysosomal membrane proteins. Although the corresponding microautophagy machinery remains to be identified, it was shown that the LC3 lipidation machinery, such as ATG5, is critical to initiate microautophagy (Lee et al., 2020). To test whether the LC3 lipidation-triggered microautophagy is involved in RNF152 degradation, we knocked out either ATG5 or ATG7 using the CRISPR-Cas9 method (Ran et al., 2013, An et al., 2019). In wildtype cells, Atg5 forms a stable 55 kDa conjugate with Atg12 in an Atg7-dependent manner (Figure 5A, left three lanes) (Mizushima et al., 1998). After knocking out Atg7, the conjugate can no longer form, and Atg5 appears as a 33 kDa band (Figure 5A, last three lanes). However, neither ATG5 nor ATG7 knockout cell lines exhibited any defect in GFP-RNF152 degradation (Figure 5A-B).

**Figure 5:**
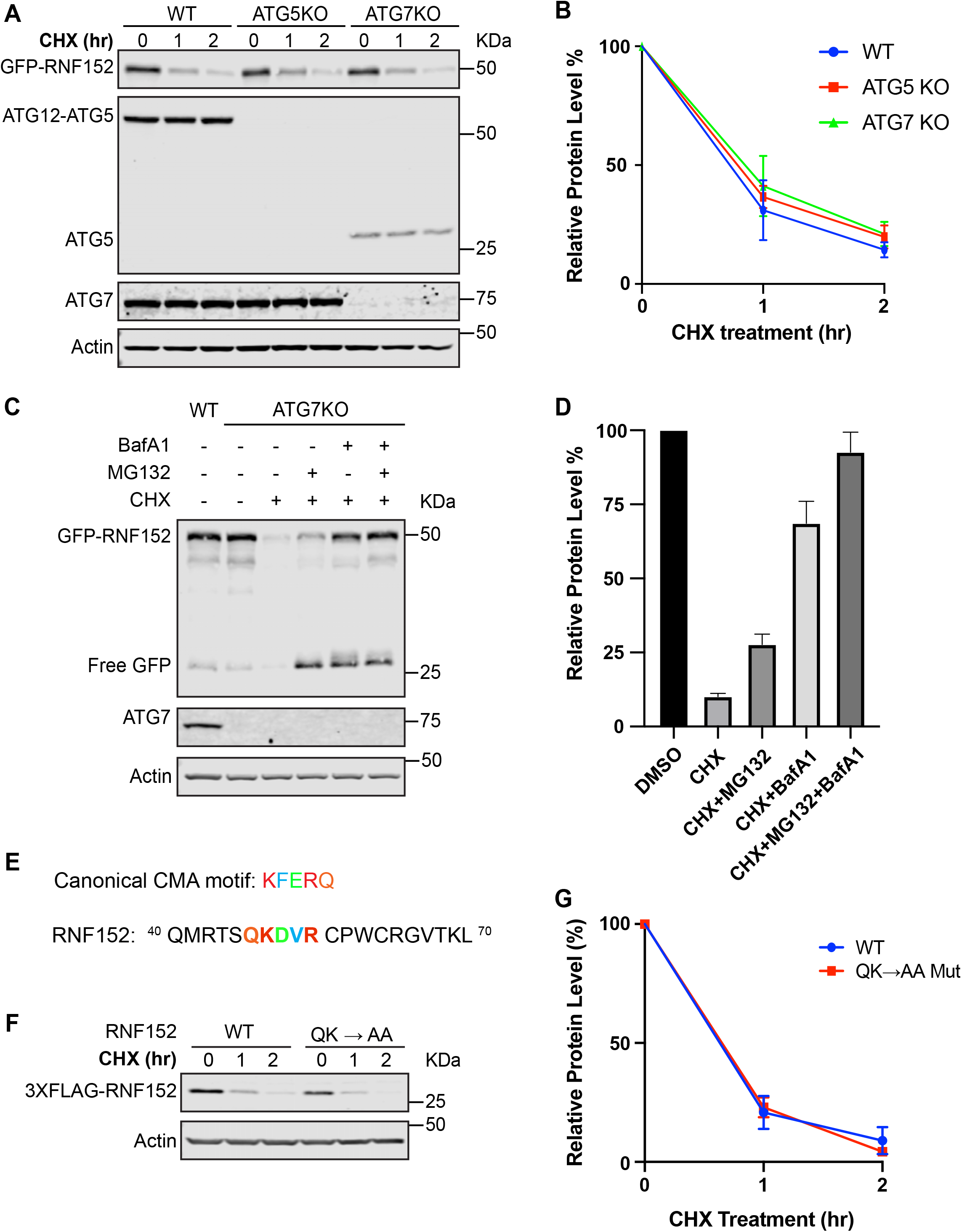
Macroautophagy and CMA pathways are not involved in the degradation of RNF152. (A) Representative Western blots showing the degradation of GFP-RNF152 in WT HEK293, ATG5KO, and ATG7KO cell lines after cycloheximide treatment. GFP-RNF152 was stably expressed in all cell lines. (B) Quantification of GFP-RNF152 level in A, n=3. (C) Representative Western blots showing GFP-RNF152 protein level in ATG7KO cells after indicated treatment for 4 hours. WT HEK293 cells serve as the control for ATG7KO. GFP-RNF152 was stably expressed in both cell lines. (D) Quantification of GFP-RNF152 level in C, n=3. (E) Identifying the canonical CMA KFERQ-like motif in RNF152 starting at residue number 46 QKDVR. (F) Representative Western blots showing the degradation of WT 3XFLAG-RNF152 and QK→AA mutant in HEK293 cells that stably express each construct after cycloheximide treatment. QK →AA: ^46^QK^47^ were mutated to alanines. (G) Quantification of F, n=3.

Autophagy and the ubiquitin-proteasome system (UPS) are the two major pathways for protein degradation in eukaryotic cells (Pohl and Dikic, 2019). There is mounting evidence that the two pathways can affect each other. It is possible that the UPS pathway is upregulated to compensate for the loss of the autophagy pathway (Wang et al., 2013; Fan et al., 2018; Kocaturk and Gozuacik, 2018; Marshall and Vierstra, 2018). To rule out the possibility that GFP-RNF152 is re-directed to the proteasome for turnover in the ATG5 or ATG7KO cell lines, we treated the ATG7KO cells that stably express GFP-RNF152 with BafA1 and/or MG132. As shown in Figure 5C-D, the degradation of full-length GFP-RNF152 is still mainly dependent on the lysosome in the macroautophagy deficient cells. Thus, the macroautophagy machinery and likely the LC3 lipidation-triggered microautophagy are not involved in RNF152 degradation.

We also examined whether chaperone-mediated autophagy (CMA) is involved. In the CMA pathway, the chaperone Hsc70 recognizes a KFERQ-like motif of a cytosolic protein and delivers it to the lysosome for degradation (Cuervo and Wong, 2014). Using a web-based “KFERQ” motif finder, we identified one putative KFERQ-like motif in the cytosolic domain of RNF152: ^46^QKDVR^50^ (Figure 5E) (Kirchner et al., 2019). Then, we mutated ^46^QK^47^ to AA and examined its degradation kinetics. As shown in Figure 5F-G, there was no significant difference in RNF152 degradation between the ^46^QK^47^→AA mutant and the WT, suggesting that the CMA pathway is not involved in the degradation of RNF152.

### The early-stage ESCRTs are less important for the degradation of GFP-RNF152

Given that macroautophagy and CMA pathways are not involved, we then asked whether ESCRT machinery is responsible for internalizing ubiquitinated GFP-RNF152. The ESCRT machinery is composed of several sub-complexes, including ESCRT-0, I, II, III, and the AAA-ATPase VPS4, all of which are conserved from yeast to human (Hurley, 2010; Henne et al., 2011). The ESCRT complex normally assembles on the surface of endosomes to sort ubiquitinated membrane proteins into the lumen as intralumenal vesicles (ILVs), creating the so-called multivesicular bodies (MVBs) (Vietri et al., 2020). Besides the ILV formation, mammalian ESCRTs play diverse roles at different membrane compartments. Recently, it was demonstrated that the ESCRT machinery could be assembled on the lysosome surface to repair damaged membranes (Radulovic et al., 2018; Skowyra et al., 2018). However, whether the ESCRT machinery can sort ubiquitinated lysosome membrane proteins such as RNF152 remains an open question. To answer this, we systematically knocked-down components of the ESCRT machinery and tested if the knockdown will affect RNF152 degradation.

We first knocked down TSG101 (ESCRT-I, Vps23 in yeast) and ALIX (accessory unit, Bro1 in yeast) using siRNA because it was shown that TSG101 and ALIX are required to recruit ESCRT-III onto the lysosome surface for membrane repair (Radulovic et al., 2018; Skowyra et al., 2018). Knocking down ALIX alone did not have an effect on RNF152 degradation. However, knocking down either TSG101 or both TSG101 and RNF152 only had a minor impact on the degradation kinetics of GFP-RNF152 (Figure S3A-B). There was also a slight increase of the ubiquitinated GFP-RNF152 (high-molecular-weight smear) in the TSG101 knockdown and TSG101+ALIX double knockdown groups (Figure S3A). This surprising result suggested that either TSG101 and ALIX might not be essential for the degradation, or a redundant component could be involved to recruit the ESCRT-III.

We also tested ESCRT-0 and ESCRT-II components by knocking down HRS (Vps27 in yeast) and VPS22, respectively. Similar to TSG101 and ALIX knockdown, HRS knockdown did not affect RNF152 degradation, but the VPS22 knockdown had a minor impact on the degradation of GFP-RNF152 (Figure S3D-F). In the case of VPS22 knockdown, we also observed a slight increase of the ubiquitinated GFP-RNF152 (Figure S3D).

To ensure the siRNA knockdown methods can truly disrupt the function of ESCRT machinery, we stained the cells with the FK2 antibody that specifically recognizes poly-ubiquitinated proteins (Firkowska et al., 2019). Since ESCRT machinery is responsible for internalizing ubiquitinated proteins from the endosome membrane, depleting ESCRT components will lead to the accumulation of poly-ubiquitinated proteins at the endosome surface. Indeed, after knocking-down either TSG101+ALIX or Vps22, we observed enlarged vacuole-like structures that stained positive with the FK2 antibody (Figure S3G). In the case of HRS knockdown, we did not observe the accumulation of poly-ubiquitinated signals on membrane structures (Figure S3G). It is unclear why this was the case. Nevertheless, the Western blot showed a pronounced reduction of the HRS protein level (Figure S3D). Besides, we noticed a severe growth defect in the HRS knockdown cells (data not shown).

Taken together, we concluded that early ESCRT components are either not essential (for ALIX and HRS) or less important (for TSG101 and VPS22) for the degradation of GFP-RNF152. Our results suggested that other redundant factors might be able to recruit the ESCRT-III to the lysosome membrane.

### The late-stage ESCRTs are important for the degradation of GFP-RNF152

We then tested the late ESCRT-III components, which polymerize into spiral filaments to drive the internalization of ubiquitinated membrane cargoes as intraluminal vesicles. After knocking down both CHMP4A and CHMP4B (Snf7 in yeast), the degradation of full-length GFP-RNF152 is significantly delayed (Figure 6A-C). Besides, we observed a substantial accumulation of ubiquitinated GFP-RNF152 in the double knockdown sample (Figure 6A). Flow Cytometry analysis further supported that knocking down both CHMP4A and CHMP4B leads to the accumulation of GFP-RNF152, as indicated by the increase of GFP fluorescence (Figure 6D). After treating cells with BafA1 to stop the lumenal degradation, we noticed an accumulation of internalized GFP-RNF152 in the siRNA control. In contrast, GFP-RNF152 appeared to accumulate on the membrane of vacuole-like LAMP2 positive structures in ~60% of cells after the double knockdown of CHMP4A and CHMP4B (Figure 6E-F).

**Figure 6:**
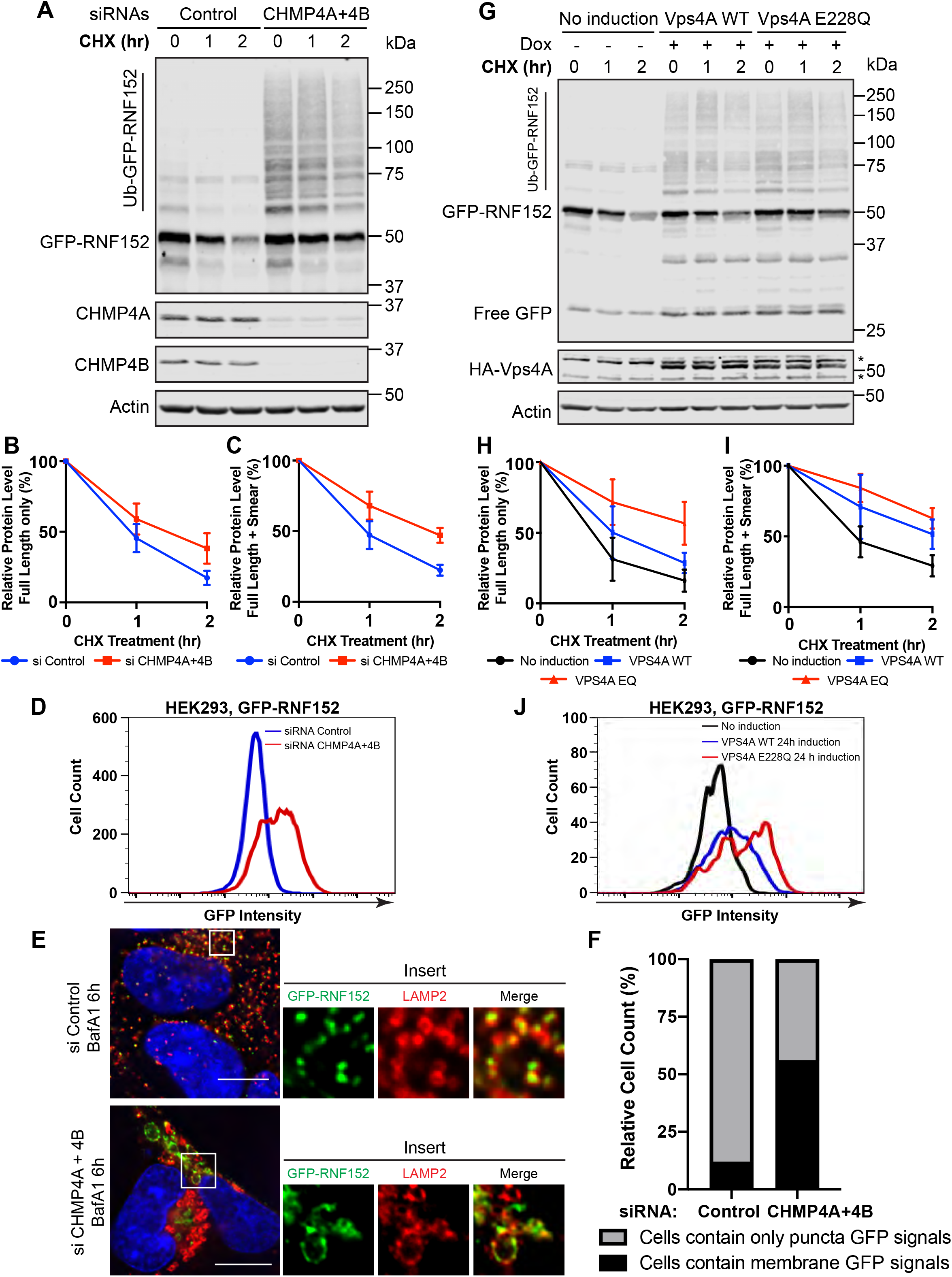
ESCRT-III and VPS4 are important for the degradation of RNF152. (A) Representative Western blots showing the cycloheximide chase of GFP-RNF152 in control and CHMP4A + CHMP4B double knockdown cells. GFP-RNF152 was stably expressed in HEK293 cells. (B) Quantification of the protein level of full-length GFP-RNF152 in A. n=3. (C) Quantification of the protein level of total GFP-RNF152 in A. n=3. (D) Flow Cytometry analysis showing GFP intensity in siRNA control and CHMP4A + CHMP4B double knockdown cells. GFP-RNF152 were stably expressed in HEK293 cells. (E) Immunofluorescence of LAMP2 with GFP-RNF152 in siRNA control and CHMP4A +CHMP4B double knockdown HEK293 cells that stably express GFP-RNF152 after 6 hours of BafA1 (400nM) treatment. Images were taken using a Delta-vision fluorescence microscopy and further deconvolved to improve the resolution. Scale bar=10 μm (F) Quantification of E showing the distribution of cells that contain either only puncta GFP signals or membrane GFP signals. (G) Representative Western blots showing the cycloheximide chase of GFP-RNF152 in cells overexpressing either WT HA-Vps4A or its dominant-negative mutant (E228Q). GFP-RNF152 were also stably expressed in these cells. To induce the expression of HA-VPS4A, cells were treated with 1μg/ml of doxycycline for 24 hours. (H) Quantification of the protein level of the full-length GFP-RNF152 at 50kDa in G, n=3. (I) Quantification of the protein level of the total GFP-RNF152 in G, n=3. (J) Flow Cytometry analysis showing GFP intensity in no induction control, WT HA-VPS4A overexpressing and HA-VPS4A E228Q overexpressing cells. GFP-RNF152 was stably expressed in these HEK293 cells.

We also validated the CHMP4A and CHMP4B knockdown by the FK2 antibody staining (Figure S4A). Double knockdown of CHMP4A and CHMP4B caused the accumulation of polyubiquitin on enlarged vacuolar structures. Together, our results suggested that the ESCRT-III components play essential roles in the degradation of GFP-RNF152.

Besides ESCRT-III, we also investigated the role of AAA-ATPase VPS4. VPS4 is responsible for the disassembly of ESCRT-III filaments. Here, we used doxycycline to induce overexpression of HA-tagged wildtype Vps4A and its dominant-negative mutant E228Q, which disrupts the AAA-ATPase activity (Takahashi et al., 2018). Overexpression of the dominant-negative mutant (E228Q) significantly delayed the degradation of full-length GFP-RNF152 (Figure 6G-I, red curve vs. black curve). Interestingly, after overexpression of wildtype HA-VPS4A, the turnover of RNF152 was also slightly delayed even though the effect was less severe than the E228Q mutant (Figure 6G-I). For both WT and E228Q mutant, we observed the accumulation of poly-ubiquitinated GFP-RNF152 (Figure 6G). These results suggested that 1) VPS4 plays a critical role in GFP-RNF152 degradation, and 2) the overexpression of HA-tagged wildtype VPS4A may also disrupt ESCRT function.

To validate our conclusions, we stained the cells with the FK2 antibody. After overexpression, both WT and the E228Q mutant showed an accumulation of FK2 positive signals, and the E228Q mutant appeared to have a much stronger effect (Figure S4B). Specifically, after overexpressing HA-VPS4A, ~50% of cells displayed FK2 positive signals, which are mainly localized to enlarged vacuolar structures. In contrast, after overexpressing the E228Q mutant, more than 90% of cells were stained positive with the FK2 antibody, and the signals were much stronger. These results indicated that overexpression of both wildtype HA-VPS4A and the dominant-negative mutant E228Q would disrupt the function of ESCRT machinery, likely due to the presence of the HA tag.

We further confirmed our finding that VPS4A plays an important role in the turnover of RNF152 using Flow Cytometry analysis. As shown in Figure 6J, overexpression of both wildtype HA-VPS4A and its dominant-negative mutant lead to an increase in the GFP intensity, indicating the stabilization of GFP-RNF152.

Taken together, our results strongly suggested that the late-stage ESCRT machinery, specifically ESCRT-III and Vps4, are important for the degradation of GFP-RNF152.

### A conserved pathway to degrade RNF152

Fundamental biological processes are generally conserved from yeast to humans. For example, the mTOR signaling complex can be found in both yeast and humans to regulate cell growth based on different nutrient levels. Here, our study uncovered a ubiquitin- and ESCRT-dependent pathway in humans to degrade lysosome membrane proteins. Is this pathway conserved in yeast? To directly answer the question, we expressed GFP-RNF152 in budding yeast. To our satisfaction, GFP-RNF152 was still correctly localized to the yeast vacuole membrane (Figure 7). Strikingly, when treated with cycloheximide, GFP-RNF152 was quickly internalized into the lumen and degraded (Figure 7A-B). Deletion of *PEP4*, the master vacuolar protease that is critical for activating other lumenal proteases, stopped the degradation of GFP-RNF152 (Figure 7A-B) (Ammerer et al., 1986; Woolford et al., 1986).

**Figure 7:**
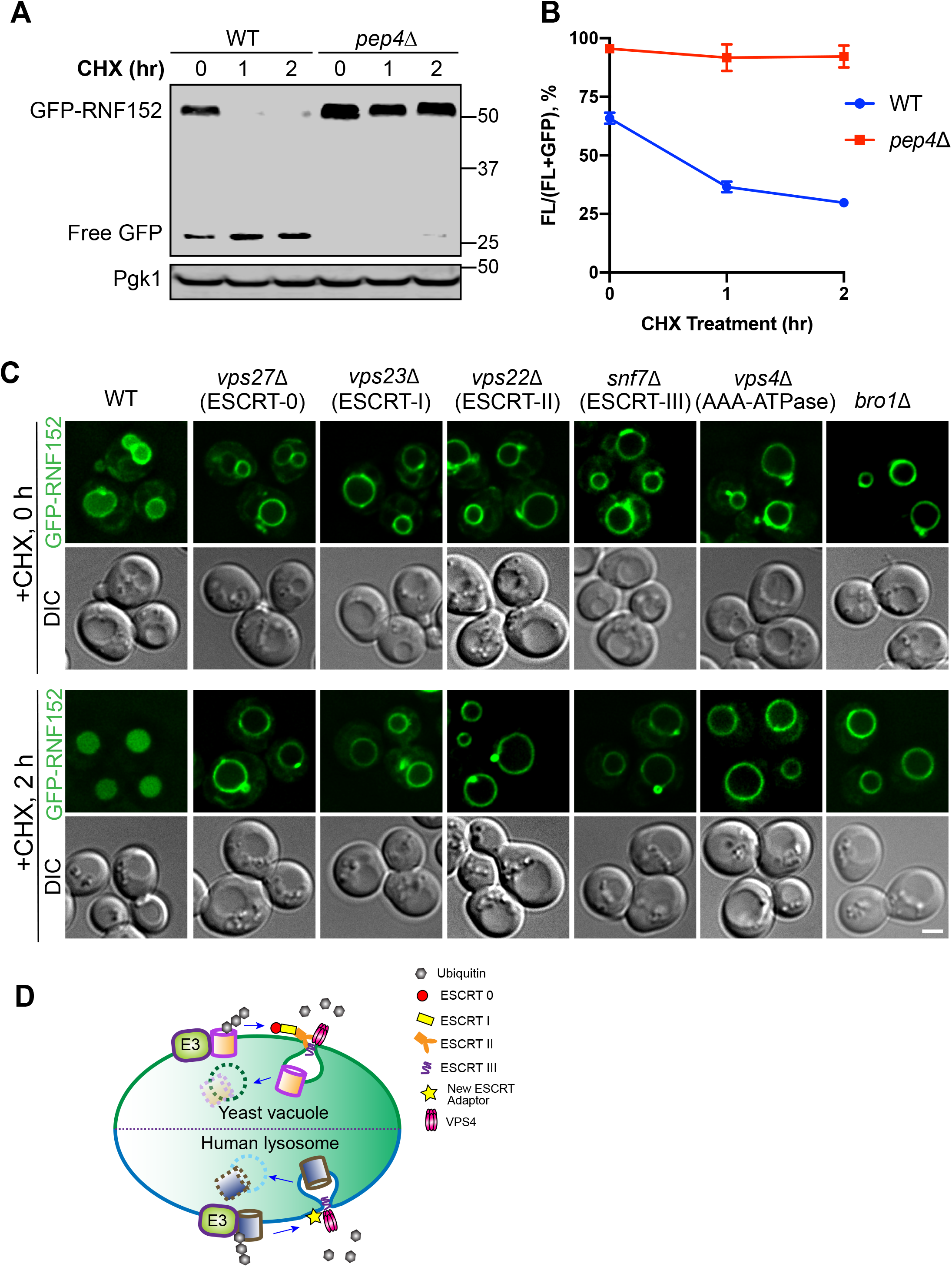
The degradation of RNF152 is ESCRT-dependent in budding yeast. (A) Representative Western blots showing the degradation of GFP-RNF152 in WT and *pep4Δ* yeast strains. (B) Quantification of the protein level in A, n=3. (C) Subcellular localization of GFP-RNF152 in WT, *vps27Δ vps23Δ vps22Δ snf7Δ vps4Δ* and *bro1Δ* yeast strains before (0h) and after (2h) cycloheximide treatment. Scale bar=2μm. DIC: differential interference contrast. (D) A Model shows the conserved ubiquitin- and ESCRT-dependent mechanism to degrade LMPs in yeast and human cells.

We then systematically deleted components of the ESCRT subcomplexes, including *VPS27* (ESCRT-0), *VPS23* (ESCRT-I), *VPS22* (ESCRT-II), *SNF7* (ESCRT-III), *VPS4* (AAA-ATPase), and *BRO1* (ALIX homolog). Because the pH (~5.5-6.5) in yeast vacuole is not low enough to quench the GFP fluorescence, we can directly observe the accumulation of free GFP during protein degradation. As shown in figure 7C, deleting any ESCRT components stabilized GFP-RNF152 on the vacuole membrane. These results indicated that the ESCRT machinery is critical for the internalization and degradation of GFP-RNF152 in yeast. Unlike human cells, early-stage ESCRT components are just as crucial as the late-stage ESCRT components in yeast, suggesting that humans may have evolved an alternative mechanism to recruit late-stage ESCRTs onto the lysosome membrane.

In summary, our investigation indicated that the ubiquitin- and ESCRT-dependent turnover of lysosome membrane proteins is a fundamental process conserved from yeast to humans (Figure 7D).

## DISCUSSION

### Different ESCRT requirements between yeast and human lysosomes

In this study, we uncovered a ubiquitin and ESCRT-dependent pathway to down-regulate human lysosome membrane proteins such as RNF152. Strikingly, when expressed in budding yeast, RNF152 is also degraded by a similar mechanism (Figure 7). Previously, we and others have demonstrated that this ubiquitin and ESCRT dependent pathway is used by budding yeast to regulate many vacuole (functional equivalent to mammalian lysosomes) membrane proteins. For example, the Rsp5-Ssh4 E3 ligase complex ubiquitinates Ypq1 (a vacuolar lysine transporter) when lysine is depleted from the cytosol (Li et al., 2015a). Similarly, a vacuolar Zn2+ influx transporter Cot1 is also regulated through ubiquitination by another E3 ligase complex, DSC, and degraded inside the vacuole (Li et al., 2015b,; Yang et al., 2018). Importantly, we and others also showed that ubiquitinated vacuole membrane proteins are internalized for degradation by the ESCRT machinery (Li et al., 2015a; Li et al., 2015b; Oku et al., 2017; Zhu et al., 2017; Morshed et al., 2020; Yang et al., 2020).

Although the ubiquitin- and ESCRT-dependent pathway is conserved from yeast to humans, we also noticed significant differences between the two systems. In budding yeast, knocking out every core component of the ESCRT machinery (ESCRT-0, I, II, III, Vps4, and BroI) stabilizes RNF152 on the vacuole membrane (Figure 7A-C). But in human cells, only ESCRT-III (CHMP4A and CHMP4B) and Vps4 AAA-ATPase activity are shown to be important to the degradation of RNF152 (Figure 6). In contrast, the early ESCRT components (ex: HRS[ESCRT-0], TSG101 [ESCRT-I], and Vps22 [ESCRT-II]) seems to be less critical (Figure S3). Our results are consistent with a previous study showing that the starvation-induced turnover of autophagy receptors by endosomal microautophagy depends on ESCRT-III and Vps4, but not the early ESCRTs (ESCRT-0, −I, and −II) (Mejlvang et al., 2018).

One possible explanation could be the functional redundancy among early components. Interestingly, many early ESCRT components share functional domains or binding features. For example, both HRS (ESCRT-0) and VPS28 (ESCRT-II) contains PI3P binding domains (Christ et al., 2017). Also, many ESCRT components, including HRS, STAM1&2 (ESCRT-0), TSG101, MVB12A&B (ESCRT-I), Vps36 (ESCRT-II), and ALIX, are all ubiquitin-binding proteins (Haglund and Dikic, 2012). Therefore, it is possible that some early ESCRT components are redundant in the degradation of LMPs.

Alternatively, it is also possible that humans may have evolved other mechanisms to recruit late-stage ESCRTs onto the lysosome membrane. During the formation of ILVs, ESCRT subcomplexes are sequentially assembled on the endosome membrane to sort and internalize ubiquitinated cargo proteins. After cargo internalization, Vps4 is recruited to disassemble and recycle the ESCRT-III sub-complex. Besides the ILV formation, ESCRTs have evolved a variety of functions on different membrane compartments in mammalian cells, such as membrane repair, budding of viral particles from the plasma membrane, midbody formation during cytokinesis, and closure of the autophagosomes. All these functions require ESCRT-III and VPS4, which are directly responsible for membrane deformation (Vietri et al., 2020). However, the corresponding proteins to recruit/assemble ESCRT-III at different membranes are quite different. For example, ESCRT-0, CEP55, and viral Gag proteins can recruit TSG101 or ALIX to distinct membrane surfaces (Vietri et al., 2020).

Many studies have suggested two parallel pathways of TSG101 (ESCRT-I) and ALIX (accessory unit of ESCRT) to independently recruit ESCRT-III to different membrane compartments. (Fujii et al., 2009; Jimenez et al., 2014; Christ et al., 2016; Tang et al., 2016; Larios et al., 2020). Here we observed that even knocking down both TSG101 and ALIX can only mildly affect the degradation of RNF152 (Figure S3A-C). These results suggested that additional factors might be involved in the degradation process. Identifying these recruitment factors will require further investigations.

### Multiple pathways may be involved in the selective turnover of LMPs

Besides the ESCRT-dependent degradation pathway examined in this study, recent studies have suggested other mechanisms might also be involved in the degradation of LMPs. First, Overholzer and colleagues showed that LC3 lipidation-triggered microautophagy is responsible for the selective turnover of several LMPs, including TRPML1 and SNAT7 (SLC38A7) (Lee et al., 2020). However, there are still many unanswered questions about this pathway. For example, how does ATG5-dependent microautophagy achieve its selectivity? It is unclear what machinery is responsible for selecting cargo proteins and how are selected LMPs sorted into the microautophagy. Moreover, how does the LC3 lipidation at the lysosome surface trigger microautophagy? Despite all these open questions, ATG5-dependent microautophagy provides an exciting model for the selective turnover of LMPs. Although our study did not find evidence that LC3 lipidation-dependent microautophagy is responsible for the degradation of RNF152, it is still possible that these two pathways may operate in parallel to regulate different subsets of LMPs.

In addition to the ESCRT-dependent pathway and LC3 lipidation-dependent microautophagy, mammalian lysosome may have evolved other mechanisms to turn over its membrane proteins, such as chaperone-mediated autophagy (CMA) and proteasome-dependent degradation. The CMA pathway often targets cytosolic proteins that contain the KFERQ-like motif. However, many membrane proteins also have the KFERQ-like motif, which potentially can be recognized by chaperone Hsc70. Therefore, It is still possible that CMA is involved in regulating some LMPs even though RNF152 is not a CMA cargo. Besides, proteasome-dependent degradation is another well-characterized and conserved pathway to down-regulate membrane proteins. The best-known example is the ER-associated degradation (ERAD) pathway, in which misfolded proteins were ubiquitinated by the E3 ligases at the endoplasmic reticulum, extracted by an AAA ATPase P97 (CDC48 in yeast), and sent to the proteasome for degradation. Interestingly, several recent studies showed that p97 could be recruited to lysosomes to facilitate the lysophagy when lysosomes are severely damaged (Papadopoulos et al., 2017, Koerver et al., 2019). Although proteasome is not directly involved in the lysophagy, it is still possible that proteasome can degrade LMP substrates extracted by p97 (Papadopoulos and Meyer, 2017). As we are still at the early stage to understand the turnover mechanisms of lysosome membrane proteins, how ESCRT machinery, autophagy factors, and proteasome may coordinate to regulate LMPs remains an exciting question to addressed by the field.

---

**Figure S1:**
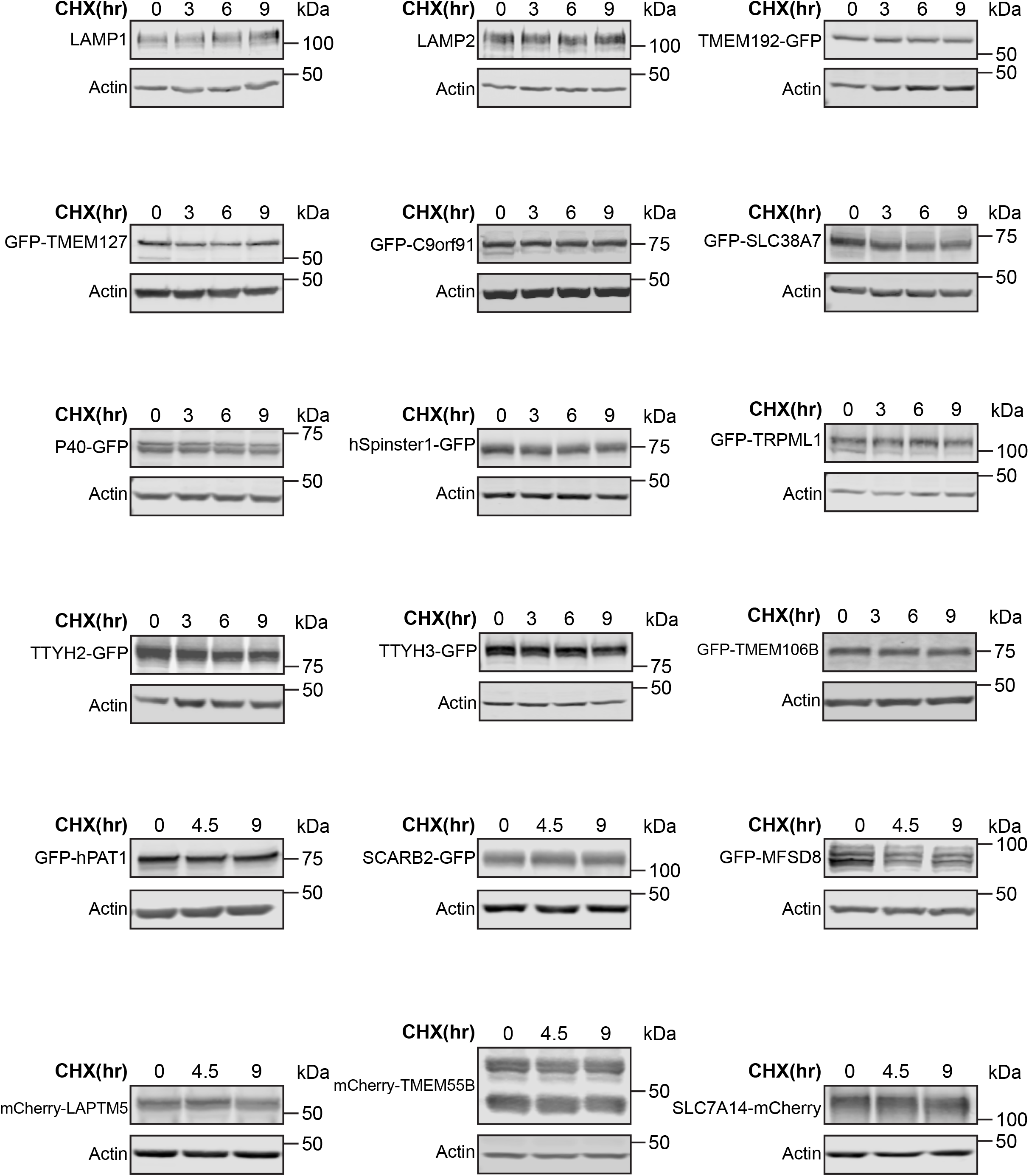
Cycloheximide chase assay for LMPs. Western blots showing examples of the CHX chase screen in HEK293 cells. LAMP1 and LAMP2 were detected at the endogenous protein level. For others, HEK293 cells were transfected with indicated overexpression plasmids. CHX chase assay was performed 20-24h after transfection with either GFP or mCherry antibody. The majority of the tested LMPs are very stable.

**Figure S2:**
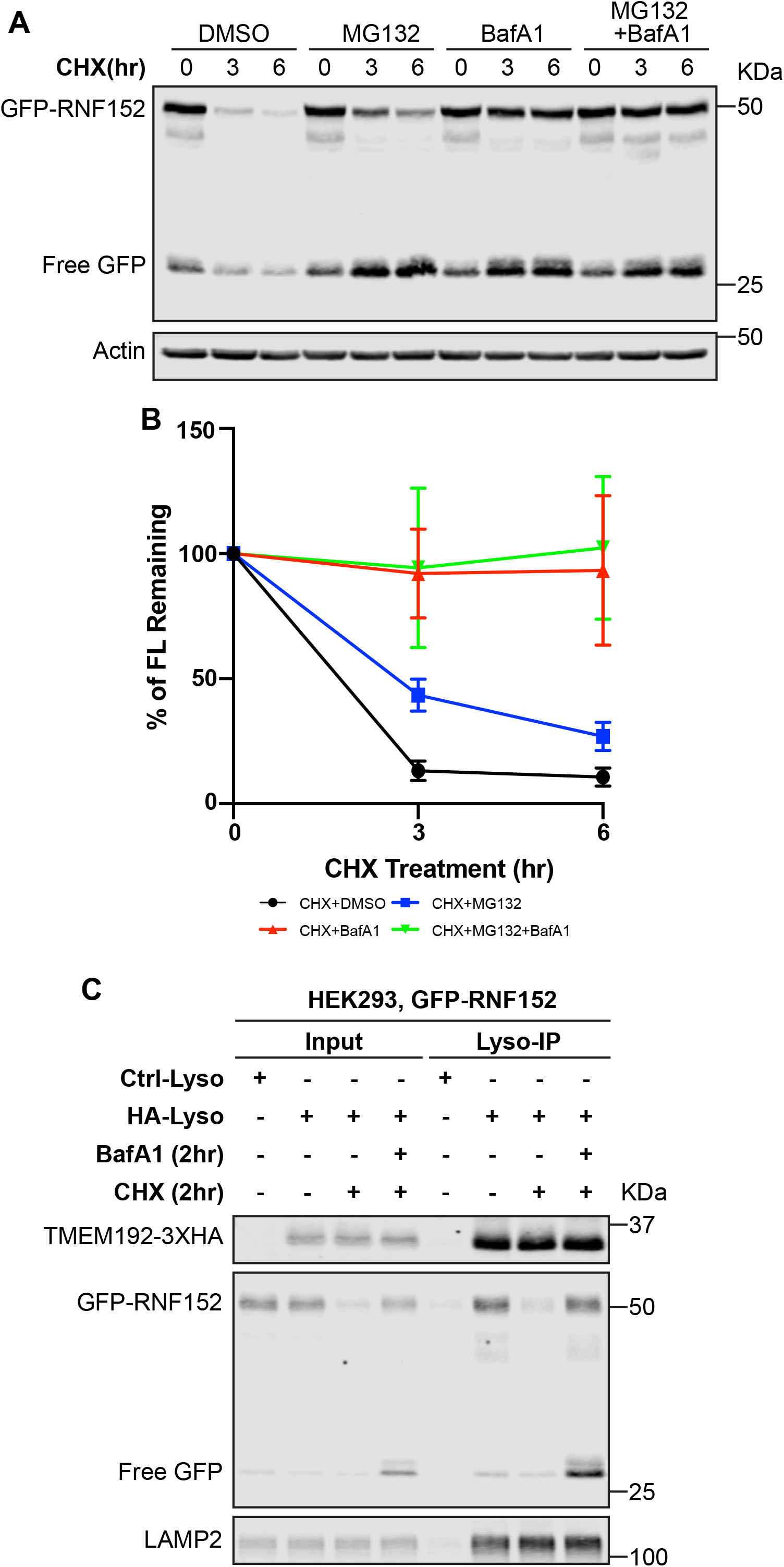
GFP-RNF152 is mainly degraded by the lysosome-dependent proteolysis. (A) Representative Western blots showing the cycloheximide chase of GFP-RNF152 with MG132 (50μM), BafA1 (400nM), and double treatment. GFP-RNF152 was stably expressed in HEK293 cells. (B) Quantification of the protein level of the full-length GFP-RNF152, n=3. (C) Representative Western blots showing that the degradation of lysosome localized GFP-RNF152 (50kDa) is blocked by BafA1 (400nM) treatment after Lyso-IP. LAMP2 serves as an internal control. Ctrl-Lyso: cells stably express TMEM192-2XFLAG. HA-Lyso: cells stable express TMEM192-3HA. Both cell lines also stably express GFP-RNF152.

**Figure S3:**
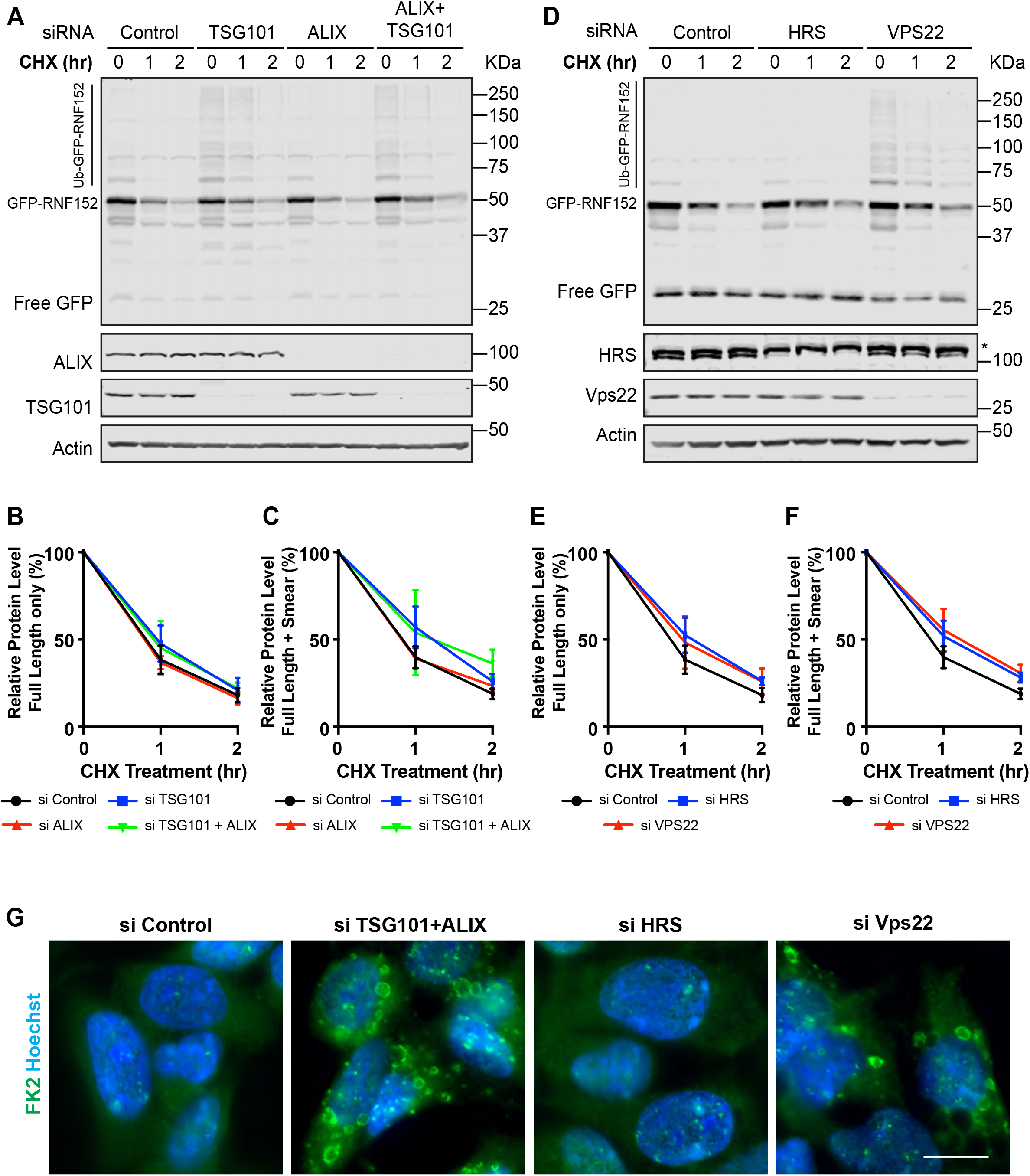
Early ESCRT components are less important for the degradation of RNF152. (A) Representative Western blots showing the cycloheximide chase of GFP-RNF152 in siRNA control, TSG101 knockdown, ALIX knockdown, and double knockdown cells. GFP-RNF152 was stably expressed in HEK293 cells. (B) Quantification of the protein level of full-length GFP-RNF152 in A, n=3. (C) Quantification of the protein level of the total GFP-RNF152 (signal at and above 50kDa) in A, n=3. (D) Representative Western blot showing the cycloheximide chase of GFP-RNF152 in siRNA control, HRS knockdown, and Vps22 knockdown cells. GFP-RNF152 was stably expressed in HEK293 cells. (E) Quantification of the full-length GFP-RNF152 in D, n=3. (F) Quantification of the protein level of the total GFP-RNF152 in D, n=3. (G) Immunostaining of poly-ubiquitinated proteins using the FK2 antibody in siRNA control HEK293 cells, HRS knockdown, TSG101+ALIX knockdown, and Vps22 knockdown cells. Nuclei were stained with Hoechst. Scale bar=10μm.

**Figure S4:**
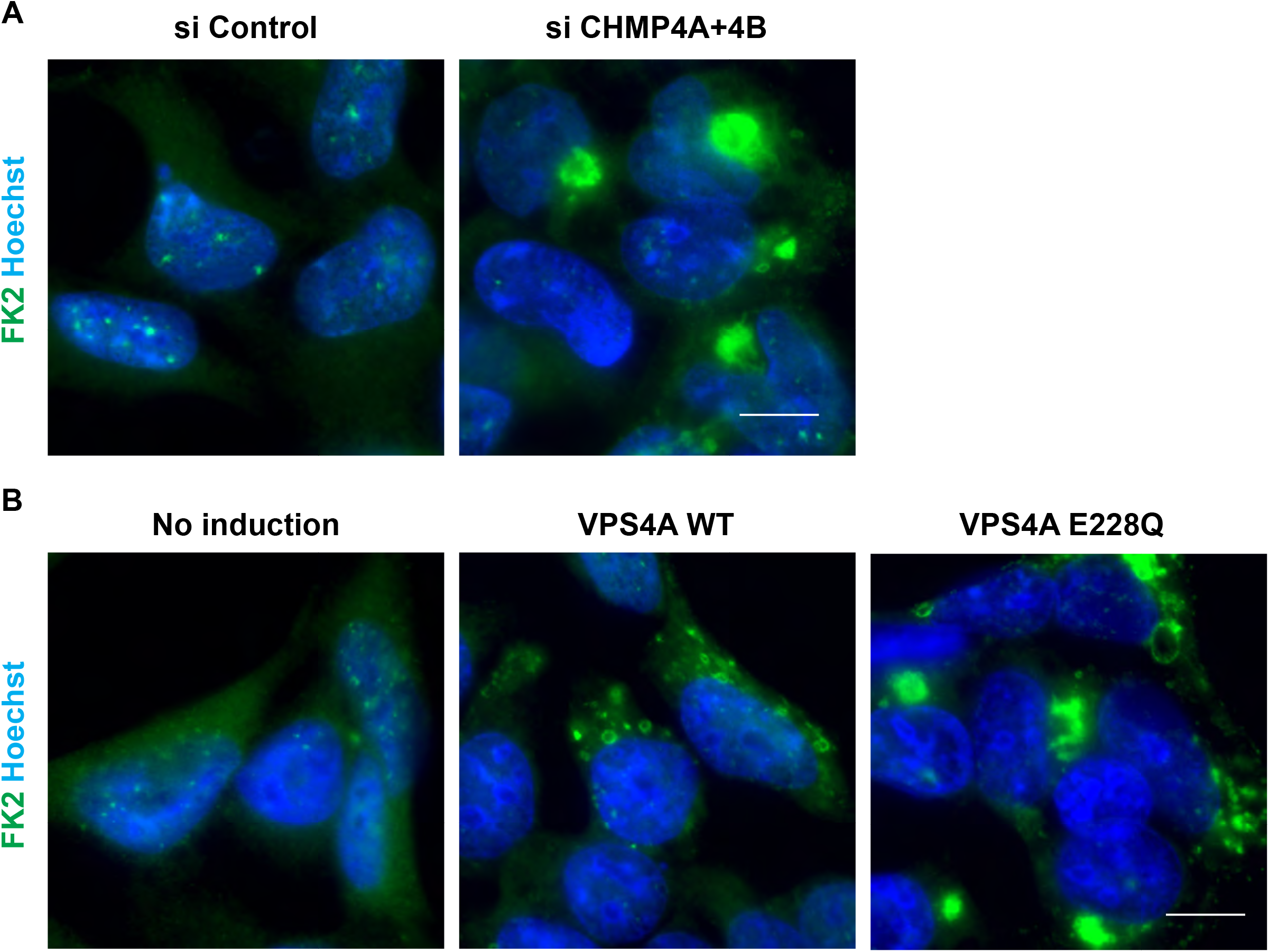
ESCRT-III knockdown and overexpression of VPS4A disrupt ESCRT function. (A) Immunostaining of poly-ubiquitinated proteins using the FK2 antibody in control HEK293 cells and CHMP4A+CHMP4B double knockdown cells. Nuclei were stained with Hoechst. Scale bar=10μm. (B) Immunostaining of poly-ubiquitinated proteins using FK2 antibody in no doxycycline induction control HEK293 cells, WT HA-Vps4A overexpressing, and HA-Vps4A E228Q overexpressing cells. The expression of HA-Vps4A was induced by 1μg/ml of doxycycline treatment for 24 hours. Scale bar=10μm

## MATERIALS AND METHODS

### Mammalian cell culture

All mammalian cell lines used in this study are listed in Table S1. The following cell lines were obtained from ATCC: HEK293 (CRL-1573), HEK293T (CRL-3216) and, HeLa (CCL-2). Cells were cultured in DMEM (Invitrogen) containing 10% Super Calf Serum (Gemini), 1% penicillin and streptomycin (Invitrogen) and 1μg/ml plasmocin (Invivogen) at 37°C, 5% CO_2_. All cells were tested negative for mycoplasma contamination using Mycoalert™ mycoplasma detection kit (Lonza).

### Plasmids

All mammalian plasmids used in this study are listed in Table S2. Most of the lysosomal membrane protein overexpression plasmids were purchased from GeneCopoeia. The CDS of RNF152 was purchased from Horizon Discovery. The 4C→S and QK→AA mutants of RNF152 were generated by PCR-based site-directed mutagenesis. The 8K→R mutant of RNF152 was synthesized as a gBlock™ gene fragment by Integrated DNA Technologies.

To generate transfer plasmids for GFP-RNF152 stable cell lines, EGFP was fused to the N-terminal of RNF152 (WT or mutants) using PCR-based overlapping extension, with 2X Gly-Gly-Gly-Ser linker in between. Then, EGFP-RNF152 was cloned into the pHAGE2-IRES-puro vector using restriction enzymes NotI and BamHI. To generate mCherry selection transfer plasmids, mCherry was amplified from the pmCherry-N1 (Clontech) vector and cloned into a pHAGE2 vector to replace the puromycin-resistant gene using restriction enzymes NdeI and ClaI. To generate transfer plasmids for 3XFLAG-RNF152 stable cell lines, RNF152 (WT or mutants) was cloned into the pBICEP-CMV2-3XFLAG (Millipore-Sigma) vector using restriction enzymes SalI and BamHI. Then, 3XFLAG-RNF152 was cloned into pHAGE2-IRES-Puro vector using restriction enzyme NcoI and BamHI.

The CDS of Vps4A E228Q was purchased from Addgene (80351). Vps4A E228Q was cloned into the pCMV-HA (Clontech) vector using restriction enzymes EcoRI and NotI. The WT HA-Vps4A was obtained by using PCR-based site-directed mutagenesis. Then the WT HA-Vps4A and HA-Vps4A E228Q were cloned into pCW57.1 (Addgene 41393) using restriction enzymes NheI and AgeI.

### Transfection and cycloheximide chase screen

HEK293 cells were cultured in DMEM containing 10% serum only media for at least 4 days before transfection. Cells were transfected with individual overexpression plasmids (2.4 μg DNA for a 3.5cm dish) using Lipofectamine 2000 (Invitrogen) according to the manufacturer’s instruction. After 20-24 hours of transfection, cells were treated with 100 μg/ml cycloheximide (Millipore-Sigma). At indicated chase time, cell samples were collected in ice-cold 1XPBS, pelleted at 800xg for 1 minute, and stored at −80°C before subsequent Western blot analysis.

### siRNA knockdown

HEK293 cells were cultured in DMEM containing 10% serum only media for at least 4 days before siRNA knockdown. Cells were transfected with siRNAs using Lipofectamine RNAimax (Invitrogen) according to the manufacturer’s instruction. After 24 hours, cells were transfected with the same amount of siRNA again. 72 hours after the first round of transfection, cells were subjected to subsequent Western blotting, immunostaining, or Flow Cytometry analysis.

The following siRNA sequences were used in this study at indicated concentration: TSG101 (50nM): 5’-CCUCCAGUCUUAUCUCGUC-dTdT-3’ (Skowyra et al., 2018) ALIX (50nM): 5’-CCUGGAUAAUGAUGAAGGATT-dTdT-3’ (Skowyra et al., 2018) VPS22 (50nM): 5’-CUUGCAGAGGCCAAGUAUA-dTdT-3’ (Christ et al., 2016) HRS (50nM): 5’-CGACAAGAACCCACACGU-dTdT-3’ (Bache et al., 2003) CHMP4A (100nM): 5’-GGCACAAACUGACGGGACA-dTdT-3’ (Mamińska et al., 2016) CHMP4B (100nM): 5’-CGAUAAAGUUGAUGAGUUA-dTdT-3’ (Mejlvang et al., 2018) ON-TARGETplus Non-targeting Pool (50nM, DharmaconTM): 5’-UGGUUUACAUGUCGACUAA-3’, 5’-UGGUUUACAUGUUGUGUGA-3’, 5’-UGGUUUACAUGUUUUUCUGA-3’, 5’-UGGUUUACAUGUUUUCCUA-3’.

### Generation of Lentiviral stable cell lines

Stable cell lines were generated as described in Abu-Remaileh et al. 2017 with some modifications (Abu-Remaileh et al., 2017). HEK293T cells were transfected with transfer plasmid, psPAX2 (Addgene 12260), and pMD2.G (Addgene 12259) at 3.5:3.5:1 ratio using Lipofectamine 2000 according to manufacturer’s instruction. Three days after transfection, the virus-containing supernatant was collected using a 5ml syringe and applied through a 0.45 μm filter. To generate stable cell lines, HEK293 or HeLa cells were seeded in 3.5cm dishes and infected with the infectious media (DMEM containing 10% super calf serum, 10 μg/ml polybrene, MOI between 0.3 to 0.4). For puromycin selection, the media was refreshed with DMEM containing 10% super calf serum and 1 μg/ml puromycin. The selection lasted for at least 7 days before subsequent analysis. For mCherry selection cells (pHAGE2-GFP-RNF152-IRES-mCherry), the media was refreshed with DMEM containing 10% super calf serum. Three days after infection, cells with proportional GFP vs. mCherry expression were enriched by FACS. Sorting was repeated 7 days after the first round of FACS.

### Generation of CRISPR-Cas9 KO cell lines

ATG5 and ATG7 knockout HEK293 cells were generated as described (Ran et al., 2013). The sgRNA guides for ATG5 and ATG7 CRISPR-Cas9 knockout were described in An et al., 2019: 5’-GATCACAAGCAACTCTGGAT-3’ for ATG5, and 5’-ATCCAAGGCACTACTAAAAG-3’ for ATG7 (An et al., 2019). In brief, sgRNA guides were ligated into pspCas9(BB)-2A-Puro (Addgene, 48139) plasmid. HEK293 cells were transfected with CRISPR-Cas9 knockout plasmids using Lipofectamine 2000 according to the manufacturer’s instruction. After 24 hours of transfection, cells were treated with 1μg/ml puromycin (Invitrogen) for 72 hours. Single cells were isolated into 96-well plates using limited dilution to a final concentration of 0.5 cell per well. The knockout colonies were screened by Western blot analysis using antibodies against ATG5 and/or ATG7. The KO cell lines were further verified by sequencing analysis to confirm the indels at target sites.

### Yeast strains, plasmids, media, and growth conditions

All yeast strains and plasmids used in this study are listed in Table S3. Difco Yeast Nitrogen Base (YNB) w/o amino acids were purchased from Millipore-Sigma. Yeast strains were grown in YNB at 26°C before further analysis.

### Mammalian sample preparation and Western blotting

Cells were collected in ice-cold 1XPBS, pelleted at 800xg for 1 minute, and lysed in lysis buffer (20mM Tris pH=8.0, 150mM NaCl, 1% Triton) containing 1Xprotease inhibitor cocktail (Biotool) at 4°C for 20 minutes. Cell lysates were centrifuged at 18,000 g for 15 minutes at 4°C. The protein concentration of the supernatant was measured by Bradford assay (Bio-rad) and normalized. After adding 2X urea sample buffer (150mM Tris pH 6.8, 6M Urea, 6% SDS, 40% glycerol, 100mM DTT, 0.1% Bromophenol blue), samples were heated at 65°C for 8 minutes. 20μg of each lysate was loaded and separated on 11% SDS-PAGE gel. Protein samples were transferred to nitrocellulose membrane for Western blot analysis. After incubated with primary and secondary antibodies, membranes were scanned using the Odyssey^®^ CLx imaging system (LI-COR).

The following primary antibodies were used for Western blotting in this study: rabbit anti-GFP (1:3000, TP401, Torrey Pines Biolabs), mouse anti-actin (1:5000, Proteintech), mouse anti-LAMP1 (1:1000, H4A3, DHSB), mouse anti-LAMP2 (1:1000, H4B4, DHSB), rabbit anti-CTSD (1:1000, Cell Signaling Technology), mouse anti-PDI (1:2000, BD Biosciences), rabbit anti-Golgin160 (1:1000, Proteintech), mouse-anti-EEA (1:500, G-4, Santa Cruz Biotechnology), rabbit-GAPDH (1:2000, Proteintech), rabbit anti-ATG5 (1:2000, D5FF5U, Cell Signaling Technology), rabbit anti-ATG7 (1:2000, D12B11, Cell Signaling Technology), rabbit anti-FLAG (1:2000, Millipore-Sigma), mouse anti-TSG101 (1:200, C-2, Santa Cruz Biotechnology), mouse anti-ALIX (1:500, 1A12, Santa Cruz Biotechnology), mouse anti-Vps22 (1:500, C-11, Santa Cruz Biotechnology), mouse anti-HRS (1:500, C-7, Santa Cruz Biotechnology), mouse anti-HA (1:1000, 16B12, BioLegend), rabbit anti-CHMP4B (1:2000, Proteintech).

The following secondary antibodies were used in this study: goat anti-mouse IRDye^®^680LT, goat anti-mouse IRDye^®^800CW, goat anti-rabbit IRDye^®^680LT, goat anti-rabbit IRDye^®^800CW. All secondary antibodies were purchased from LI-COR Biosciences and used at 1:10,000 dilution.

### Yeast sample preparation and Western blotting

Typically, 7 OD_600_ unit yeast cells were collected for sample preparation at each timepoint. The cells were first resuspended with 10% ice-cold TCA and incubated on ice for at least 1 hr. After washing with 0.1% TCA, the cell pellets were then resuspended in 70 μl 2× boiling buffer (150 mM Tris, pH 6.8; 6 M urea; 6% SDS; 10% glycerol; 100 mM DTT), lysed by bead beating for 5 minutes, and heat-treated at 65°C for another 5 minutes. After the addition of 70 μl 2 × urea sample buffer (50 mM Tris, pH 7.5; 6 M urea; 10 mM EDTA; 2% SDS; 100 mM DTT, and 0.1% bromophenol blue), samples were treated for another round of bead beating and 65°C heating for 5 minutes, respectively. After spinning at 13,000 g for 5 minutes, the supernatant was collected, subjected to SDS-PAGE, and transferred to nitrocellulose membranes for Western blotting analysis. One OD_600_ unit (20 μl) of yeast cells was loaded in each lane.

The following primary antibodies were used in this study: mouse anti-Pgk1 (1:5,000, 22C5D8; Invitrogen) and rabbit anti-GFP (1:3000, TP401, Torrey Pines Biolabs).

### Immunostaining for mammalian samples

All incubation processes were performed in dark. Cells grown on 1.5 circular glass coverslips were washed with ice-cold 1XPBS and fixed in cold 100% methanol for 8 minutes at −20°C. The fixed samples were blocked in 3% BSA (in 1XPBS) for 30 minutes at room temperature followed by incubating with primary and secondary antibodies. The Cell Nucleus was stained using Hoechst (1:8000, Invitrogen). Coverslips were mounted in Fluoromount-G^®^ (SouthernBiotech) and cured for at least 24 hours before imaging.

Samples were imaged with either a DeltaVision fluorescence microscope (GE Healthcare Life Science) or a Leica SP8 confocal microscopy. For images taken with the DeltaVision fluorescence microscope, deconvolution was performed with the softWoRx program. Images were further cropped and adjusted using ImageJ (National Institutes of Health).

The following primary antibodies were used for immunostaining in this study: mouse anti-LAMP2 (1:100, H4B4, DHSB), mouse anti-ubiquitin (1:100, FK2, Millipore-Sigma).

The following secondary antibodies were used in this study: FITC goat anti-mouse (1:100, Jackson ImmunoReseach) and TRITC goat anti-mouse (1:100, Jackson ImmunoResearch).

### Yeast Microscopy and image processing

Yeast cells were collected and washed with MilliQ water once before imaging. The microscopy and imaging processing were performed with a DeltaVision system (GE Healthcare Life Sciences) as described in Yang et al., 2018, and the FITC (excitation, 475/28; emission, 525/48) filter set was used for GFP channel. Image acquisition and deconvolution were performed with the softWoRx program. The images were further cropped or adjusted using ImageJ (National Institutes of Health).

### Ubiquitin immunoprecipitation with GFP-Trap^®^A nanobody

The HA-Ubiquitin plasmid was transfected into HEK293 cells that either stably express free GFP or GFP-RNF152. Immunoprecipitation was performed 48 hours post-transfection according to the manufacturer’s instruction with some modifications. In brief, cells (one 10 cm dish of near-confluent cells per IP group) were collected in ice-cold 1XPBS, pelleted at 1000 g for 1 minute, and lysed in 300 μl of lysis buffer (20mM Tris pH=8.0, 150mM NaCl, 1% Triton) containing 100 μM of N-Ethylmaleimide (Millipore-Sigma) and 1Xprotease inhibitor cocktail (Biotool) at 4°C for 20 minutes. Cell lysates were centrifuged at 18,000 g for 15 minutes at 4°C. The concentration of the supernatant was measured by Bradford assay (Bio-rad) and normalized. 15 μl of GFP-Trap^®^A (pre-equilibrated with lysis buffer, Chromotek) was added to 285μl of normalized cell lysate and incubated at 4°C for 2 hours with gentle rocking. The resin was then washed once with lysis buffer, three times with stringent washing buffer (8M Urea, 1%SDS in 1XPBS), and once with 1%SDS in 1XPBS. To elute bound proteins, the resin was incubated with 2X Urea sample buffer (150mM Tris pH 6.8, 6M Urea, 6% SDS, 40% glycerol, 100mM DTT, 0.1% Bromophenol blue) at 65°C for 10 minutes. The resulting eluates were analyzed by Western blotting.

### Lyso-IP

LysoIP was conducted as described before (Abu-Remaileh et al., 2017) with some modifications. About 2X107 cells in a 15cm dish were used for each LysoIP experiment. Cells were rinsed twice with cold PBS, scraped, and collected with 1ml KPBS (136 mM KCl, 10 mM KH_2_PO_4_, pH=7.25). Cells were centrifuged at 1000 g for 2 minutes at 4℃. Pelleted cells were resuspended in 1ml of KPBS with protease inhibitor and homogenized. The homogenate was then centrifuged at 1000 g for 2 minutes at 4℃. For input, 40μl of supernatant was taken (about 5% to the total amount) and mixed with 2X Urea sample buffer (150mM Tris pH 6.8, 6M Urea, 6% SDS, 40% glycerol, 100mM DTT, 0.1% Bromophenol blue). For IP, 800μl supernatant was incubated with 20μl anti-HA beads (Millipore-Sigma) and rotated for 20 minutes. The beads were washed with KPBS 6 times. To eluate bound lysosomes, the beads were resuspended in 40μl KPBS and 2X Urea sample buffer (150mM Tris pH 6.8, 6M Urea, 6% SDS, 40% glycerol, 100mM DTT, 0.1% Bromophenol blue). Samples were heated at 65°C for 10 minutes, followed by Western blot analysis.

### Flow Cytometry analysis and FACS

Cells were washed with 1XPBS and trypsinized until all cells are dissociated from the dishes. Dissociated cells were neutralized with DMEM containing 10% serum media and pelleted at 300xg for 3 minutes. Cells were resuspended in ice-cold 1XPBS and analyzed using either an LSR Fortessa (BD Biosciences) or a Ze5 (Bio-rad) flow cytometer. For FACS, cells were sorted using FACSAria III cell sorter (BD Biosciences). Flow cytometry analysis and FACS were performed by technicians from the Flow Cytometry Core at the University of Michigan. The data were analyzed using FlowJo software.

### Quantification and statistical analysis

The band intensity for Western blot was quantified using Image Studio software (LI-COR). Graphs were generated using Prism (GraphPad). Statistical analysis was performed with the Student t-test. Error bars represent the standard deviation.

---

**Supplemental Table 1:**
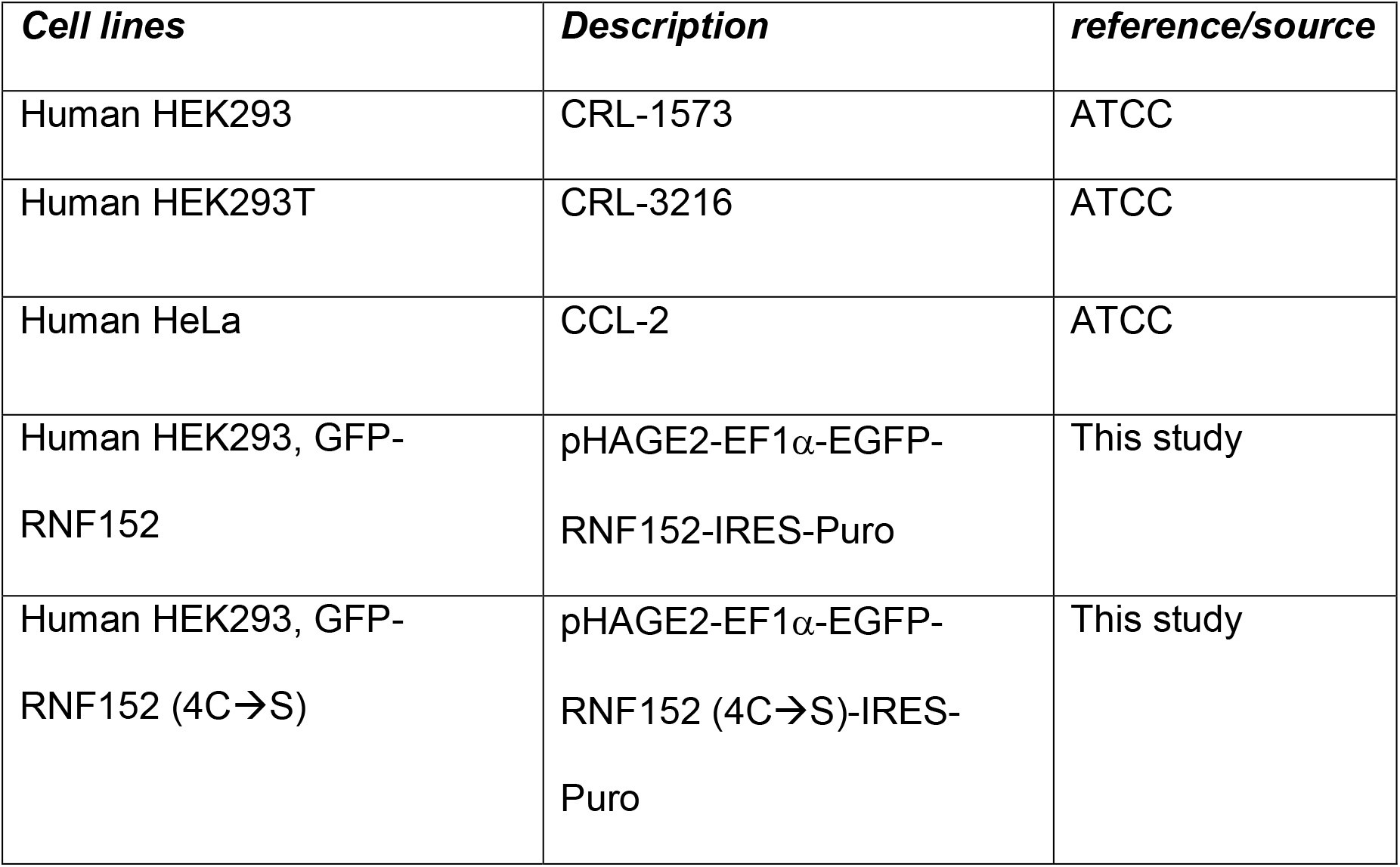
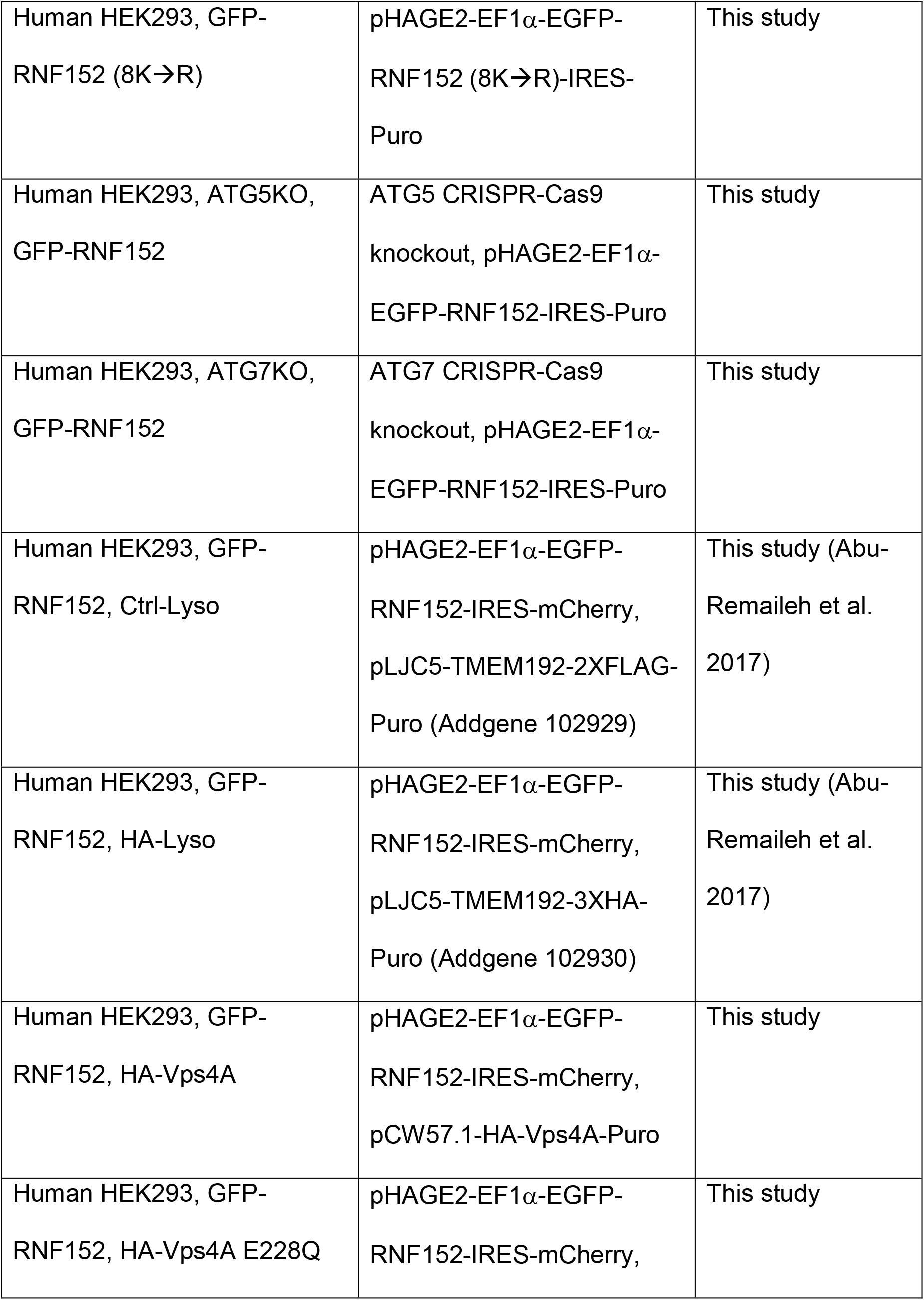
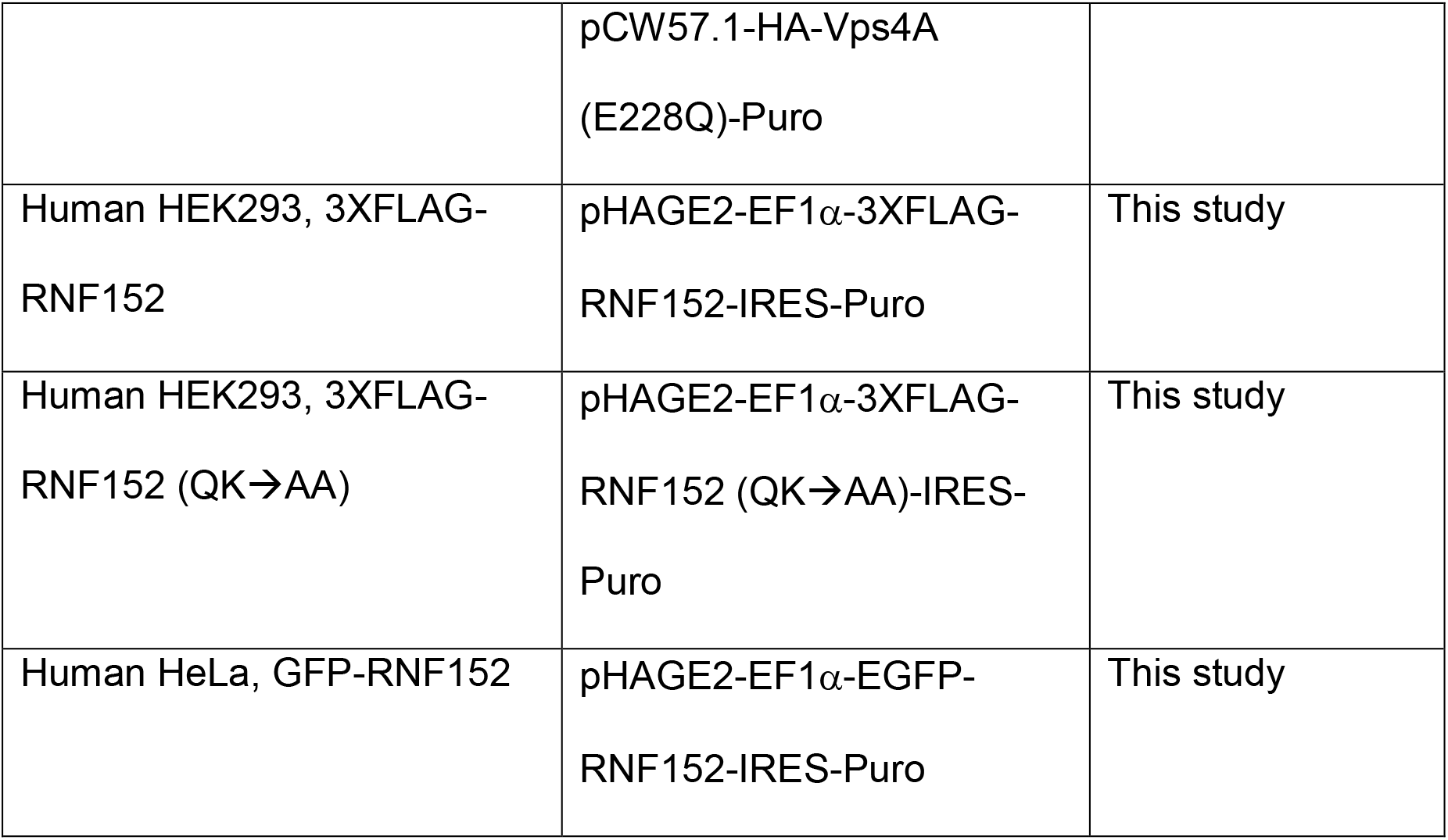
Mammalian cell lines used in this study.

| **Supplemental Table 1: Mammalian cell lines used in this study** |  |  |
| --- | --- | --- |
| ***Cell lines*** | ***Description*** | ***reference/source*** |
| Human HEK293 | CRL-1573 | ATCC |
| Human HEK293T | CRL-3216 | ATCC |
| Human HeLa | CCL-2 | ATCC |
| Human HEK293, GFP-RNF152 | pHAGE2-EF1α-EGFP-RNF152-IRES-Puro | This study |
| Human HEK293, GFP-RNF152 (4C→S) | pHAGE2-EF1α-EGFP-RNF152 (4C→S)-IRES-Puro | This study |
| Human HEK293, GFP-RNF152 (8K→R) | pHAGE2-EF1α-EGFP-RNF152 (8K→R)-IRES-Puro | This study |
| Human HEK293, ATG5KO, GFP-RNF152 | ATG5 CRISPR-Cas9 knockout, pHAGE2-EF1α-EGFP-RNF152-IRES-Puro | This study |
| Human HEK293, ATG7KO, GFP-RNF152 | ATG7 CRISPR-Cas9 knockout, pHAGE2-EF1α-EGFP-RNF152-IRES-Puro | This study |
| Human HEK293, GFP-RNF152, Ctrl-Lyso | pHAGE2-EF1α-EGFP-RNF152-IRES-mCherry, pLJC5-TMEM192-2XFLAG-Puro (Addgene 102929) | This study (Abu-Remaileh et al. 2017) |
| Human HEK293, GFP-RNF152, HA-Lyso | pHAGE2-EF1α-EGFP-RNF152-IRES-mCherry, pLJC5-TMEM192-3XHA-Puro (Addgene 102930) | This study (Abu-Remaileh et al. 2017) |
| Human HEK293, GFP-RNF152, HA-Vps4A | pHAGE2-EF1α-EGFP-RNF152-IRES-mCherry, pCW57.1-HA-Vps4A-Puro | This study |
| Human HEK293, GFP-RNF152, HA-Vps4A E228Q | pHAGE2-EF1α-EGFP-RNF152-IRES-mCherry, | This study |
|  | pCW57.1-HA-Vps4A<br>(E228Q)-Puro |  |
| Human HEK293, 3XFLAG-RNF152 | pHAGE2-EF1α-3XFLAG-RNF152-IRES-Puro | This study |
| Human HEK293, 3XFLAG-RNF152 (QK→AA) | pHAGE2-EF1α-3XFLAG-RNF152 (QK→AA)-IRES-Puro | This study |
| Human HeLa, GFP-RNF152 | pHAGE2-EF1α-EGFP-RNF152-IRES-Puro | This study |

**Supplemental Table 2:** Mammalian plasmids used in this study.

| **Supplemental Table 2: Mammalian plasmids used in this study** |  |  |  |
| --- | --- | --- | --- |
| ***Vector*** | ***Insert*** | ***description*** | ***reference/source*** |
| pEGFP-C1 | RNF152 | CMV promoter, N-terminal GFP | This study |
| pBICEP-CMV2-3XFLAG | RNF152 | CMV promoter, N-terminal 3XFLAG | This study |
| pHAGE2-IRES-Puro | EGFP | EF1α promoter, puromycin selection | This study |
| pHAGE2-IRES-Puro | EGFP-RNF152 | EF1α promoter, puromycin selection | This study |
| pHAGE2-IRES-Puro | EGFP-RNF152 (4C→S) | EF1α promoter, puromycin selection | This study |
| pHAGE2-IRES-Puro | EGFP-RNF152 (8K→R) | EF1α promoter, puromycin selection | This study |
| pHAGE2-IRES-Puro | 3XFLAG-RNF152 | EF1α promoter, puromycin selection | This study |
| pHAGE2-IRES-Puro | 3XFLAG-RNF152 (QK→AA) | EF1α promoter, puromycin selection | This study |
| pHAGE2-IRES-mCherry | EGFP-RNF152 | EF1α promoter, mCherry selection | This study |
| pEGFP-C1 | Vps4 E228Q | CMV promoter, N-terminal GFP | Votteler et al. 2016<br>Addgene 80351 |
| pCMV-HA | HA-Vps4A E228Q | CMV promoter, N-terminal HA | This study |
| pCW57.1 | HA-Vps4A WT | Tet-on promoter | This study |
| pCW57.1 | HA-Vps4A E228Q | Tet-on promoter | This study |
| pCMV-HA-Ub |  | CMV promoter, three repeats of HA-Ub | This study |
| pLJC5 | TMEM192-2XFLAG | UbC promoter | Abu-Remaileh et al. 2017<br>Addgene 102929 |
| pLJC5 | TMEM192-3XHA | UbC promoter | Abu-Remaileh et al. 2017<br>Addgene 102930 |
| pSpCas9(BB)-2A-Puro (PX459) |  | CRISPR-Cas9 knockout | Ran et al. 2013<br>Addgene, 48139 |
| psPAX2 |  | Lentiviral packaging plasmid | Addgene 12260 |
| pMD2.G |  | VSV-G envelope | Addgene 12259 |
| *Overexpression plasmids used in the cycloheximide chase screen* |  |  |  |
| pcDNA3-EGFP | CLCN7 | CMV promoter, C-terminal GFP | This study. CDS was purchased from Origene. |
| pCMV6-AC-GFP | OSTM1 | CMV promoter, C-terminal turboGFP | This study, Origene |
| pCMV-SPORT6 | p40-EGFP | CMV promoter, C-terminal GFP | Gift from Dr. M Boonen. (Boonen et al. 2006) |
| pEGFP-C1 | hPAT1<br>(SLC36A1) | CMV promoter, N-terminal GFP | This study, CDS purchased from Origene Technologies |
| pEGFP-C1 | TMEM192 | CMV promoter, N-terminal GFP | Gift from B. Schröder.<br>(Schröder et al. 2010) |
| pEGFP-C2 | TRPML1 | CMV promoter, N-terminal GFP | Dong et al. 2008. |
| pEGFP-N1 | CTNS | CMV promoter, C-terminal GFP | This study, CDS purchased from DNASU. |
| pEGFP-N1 | hSpinster1 | CMV promoter, C-terminal GFP | This study. (Rong et al. 2011) |
| pEGFP-N1 | SCARB2 | CMV promoter, C-terminal GFP | This study, CDS purchased from DNASU. |
| pReceiver-M03-EGFP | PQLC2 | CMV promoter, C-terminal GFP | This study<br>GeneCopoeia |
| pReceiver-M03-EGFP | TTYH2 | CMV promoter, C-terminal GFP | This study<br>GeneCopoeia |
| pReceiver-M03-EGFP | TTYH3 | CMV promoter, C-terminal GFP | This study<br>GeneCopoeia |
| pReceiver-M29-EGFP | C9orf91 | CMV promoter, N-terminal GFP | This study<br>GeneCopoeia |
| pReceiver-M29-EGFP | ITM2C | CMV promoter, N-terminal GFP | This study<br>GeneCopoeia |
| pReceiver-M29-EGFP | MFSD8 | CMV promoter, N-terminal GFP | This study<br>GeneCopoeia |
| pReceiver-M29-EGFP | OCA2 | CMV promoter, N-terminal GFP | This study<br>GeneCopoeia |
| pReceiver-M29-EGFP | SLC38A7 | CMV promoter, N-terminal GFP | This study<br>GeneCopoeia |
| pReceiver-M29-EGFP | STARD3 | CMV promoter, N-terminal GFP | This study<br>GeneCopoeia |
| pReceiver-M29-EGFP | TMEM106B | CMV promoter, N-terminal GFP | This study<br>GeneCopoeia |
| pReceiver-M29-EGFP | TMEM127 | CMV promoter, N-terminal GFP | This study<br>GeneCopoeia |
| pReceiver-M29-EGFP | TMEM175 | CMV promoter, N-terminal GFP | This study<br>GeneCopoeia |
| pReceiver-M55-mCherry | LAPTM5 | CMV promoter, N-terminal mCherry | This study<br>GeneCopoeia |
| pReceiver-M55-mCherry | TMEM55B | CMV promoter, N-terminal mCherry | This study<br>GeneCopoeia |
| pReceiver-M56-mCherry | LAPTM4A | CMV promoter, C-terminal mCherry | This study<br>GeneCopoeia |
| pReceiver-M56-mCherry | SLC7A14 | CMV promoter, C-terminal mCherry | This study<br>GeneCopoeia |

**Supplemental Table 3:**
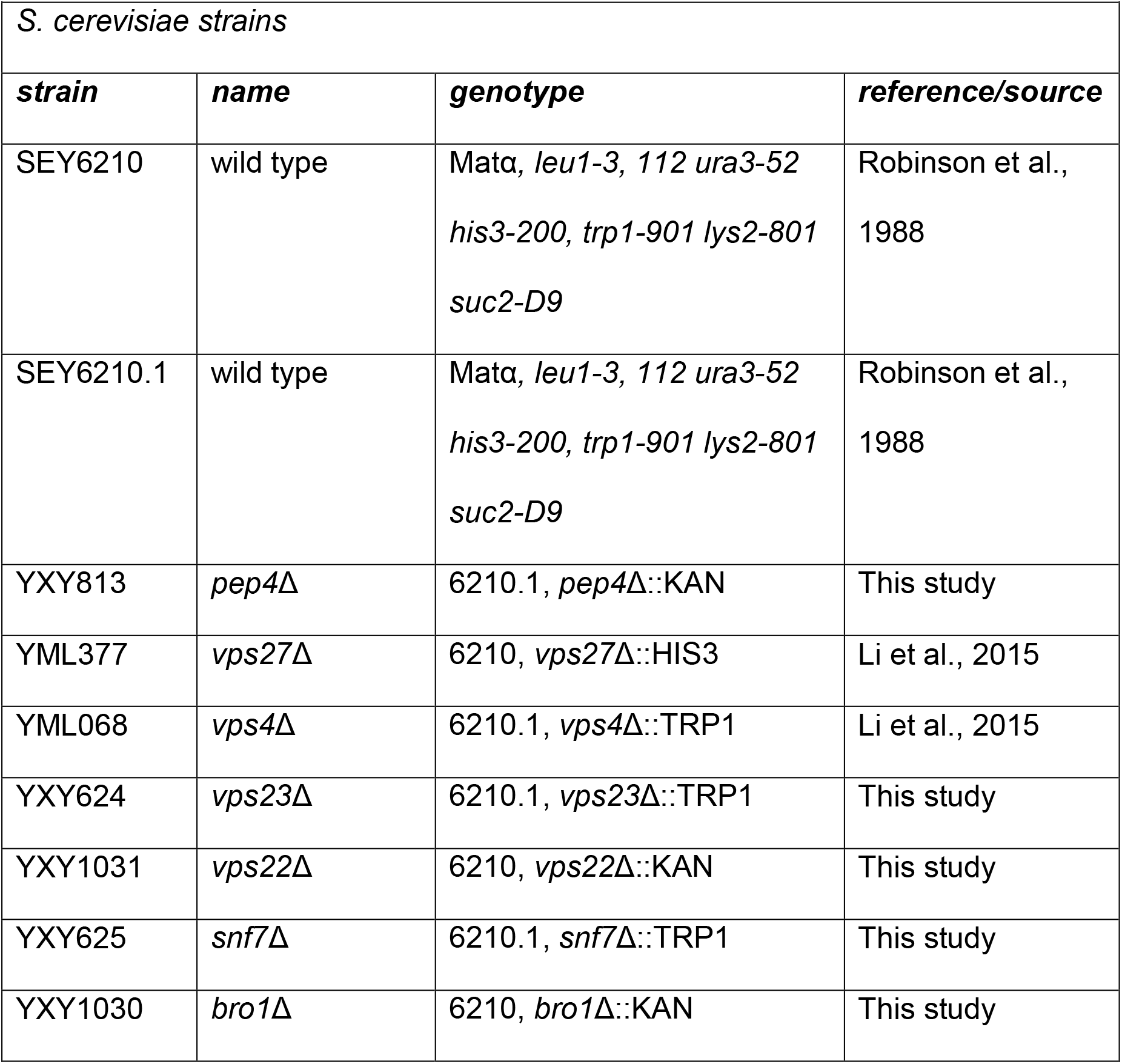
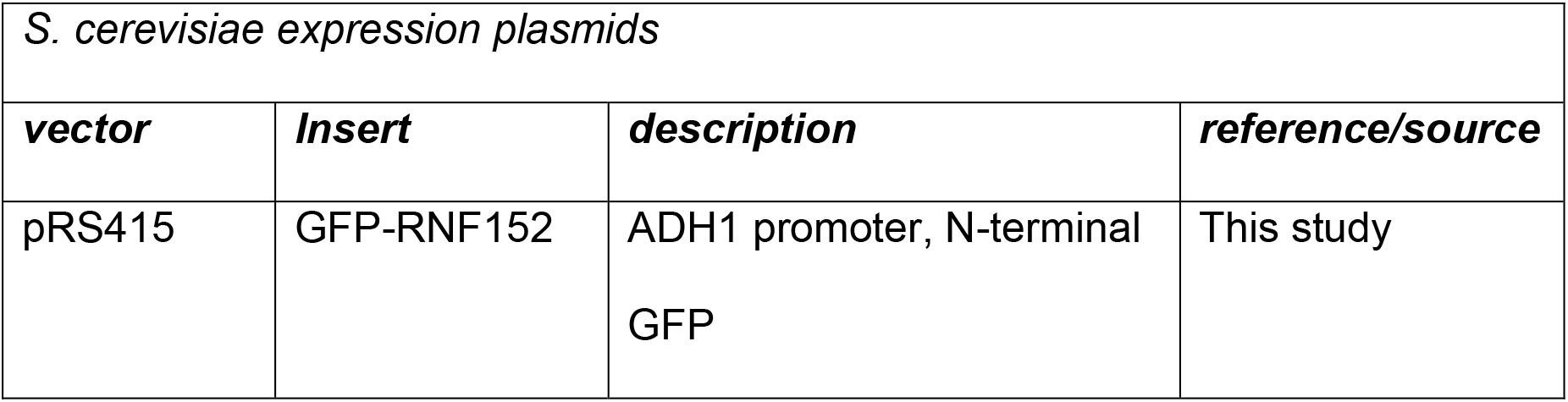
Yeast strains and Plasmids used in this study.

